# LONP-1 deficiency causes dysregulated protein synthesis within mitochondria that is restored by mitoribosomal mutations

**DOI:** 10.64898/2026.01.14.699555

**Authors:** Levi Ali, Avijit Mallick, Yunguang Du, Rui Li, Lihua Julie Zhu, Kuang Shen, Cole M. Haynes

**Affiliations:** Department of Molecular, Cell, and Cancer Biology, University of Massachusetts Chan Medical School, Worcester, MA 01605, USA

**Keywords:** LONP1, ATFS-1, MRPS38, MRPS15, mitoribosome, mitochondria, OXPHOS, mtDNA, protein synthesis

## Abstract

Mitochondrial homeostasis is maintained by multiple molecular chaperones and proteases located within the organelle. The mitochondrial matrix-localized protease LONP-1 degrades oxidatively damaged or misfolded proteins. Importantly, LONP-1 also regulates mitochondrial DNA replication. Here, we show that mutations in *C. elegans* that impair LONP-1 function cause dysregulation of mitochondrial DNA replication, mitochondrial RNA transcription and protein synthesis within the mitochondrial matrix. LONP-1 deficient worms had reduced levels of oxidative phosphorylation proteins despite increased mtDNA-encoded protein synthesis. Via a forward genetic screen, we identified three mutations that restored mitochondrial function and the rate of development in *lonp-1* mutants to levels comparable to those in wildtype worms. Interestingly, all three suppressor mutations were found in genes encoding mitochondrial ribosome proteins. A point mutation in the mitochondrial ribosome protein MRPS-38 restored oxidative phosphorylation in *lonp-1* mutant worms. Combined, our results suggest that LONP-1 regulates mitochondrial protein synthesis and that the suppressor mutations within MRPS-38 or MRPS-15 enhance oxidative phosphorylation complex assembly by slowing translation.

## Introduction

The production of ATP by oxidative phosphorylation (OXPHOS) is a central role of mitochondria. OXPHOS requires multimeric protein complexes localized within the mitochondrial inner membrane. Nearly 100 assembly factors generate the OXPHOS complexes, which together are comprised of approximately 90 proteins^1^. Importantly, 13 essential OXPHOS proteins (12 in *C. elegans*) are encoded by mitochondrial DNA (mtDNA) and synthesized on ribosomes within the mitochondrial matrix^2^. The requirement of hundreds of proteins for biogenesis of the OXPHOS complexes highlights the complexity of nuclear–mitochondrial coordination between genomes that supports this essential function.

The mechanisms by which protein homeostasis and quality control are maintained during mitochondrial biogenesis and during mitochondrial stress are not fully understood. Importantly, numerous proteins that function in translation of mtDNA-encoded proteins have been linked to inherited genetic diseases that currently lack effective treatments^3,4^. It was recently discovered that newly synthesized mtDNA-encoded proteins are actively monitored, with unassembled translation products being rapidly degraded by the membrane-embedded prohibitin (PHB)/m-AAA protease supercomplexes^5^. This finding highlights the importance of regulating nascent proteins within mitochondria and hints at a complex system of quality control dependent on multiple mitochondrial proteases.

Considerable evidence indicates that disruption of mitochondrial proteostasis promotes the formation of protein aggregates within the mitochondrial matrix, which is thought to contribute to neurodegenerative disease^6,7^. Mitochondrial protein homeostasis is maintained by molecular chaperones and proteases that facilitate protein folding, assembly, and turnover^8^. The highly conserved protease Lon (hereafter termed LONP-1) is present in archaea, bacteria and the mitochondria of eukaryotes^9^. LONP-1 recognizes substrates by interacting with hydrophobic domains and maintains protein quality control by selective degradation of oxidatively damaged or misfolded proteins^10,11^. Studies in yeast and human cells have established an additional role for LONP-1 as a molecular chaperone that promotes the assembly of OXPHOS complexes^12–15^. Deletion of the *lonp-1* gene causes embryonic lethality in flies and mice.^16,17^ In humans, missense mutations in LONP-1 can cause CODAS syndrome, a fatal condition characterized by cerebral, ocular, dental, auricular and skeletal anomalies^18,19^.

Here, we find that premature stop codon mutations in the *lonp-1* gene cause severe mitochondrial dysfunction which activates the mitochondrial unfolded protein response (UPR^mt^), resulting in increased mtDNA copy number, mtRNA-encoded transcripts and mitochondrial protein synthesis in the nematode *C. elegans*. Via a forward genetic screen, we identified 3 distinct mutations in nuclear-encoded genes that improve mitochondrial function in *lonp-1* mutant worms. Among these, a point mutation in the mitochondrial ribosomal protein MRPS-38 restores diminished OXPHOS complexes and reduced organismal development caused by OXPHOS assembly defects. Our findings highlight that LONP-1 plays a significant role in regulating mtDNA-encoded protein translation and OXPHOS assembly, and that a mutation in the mitochondrial ribosomal protein MRPS-38 can restore OXPHOS when LONP-1 is impaired.

## Results

### Mutations in *lonp-1* cause severe mitochondrial dysfunction

To gain insight into the essential functions of the conserved mitochondrial matrix-localized protease LONP-1, we generated two mutant *C. elegans* strains using CRISPR-cas9 genome editing. The *lonp-1(cmh24)* allele contains a stop cassette in the first exon, while *lonp-1(cmh25)* carries a premature stop codon in the fifth exon (Figure 1A). Western blot analysis shows no detectable LONP-1 protein in either mutant, consistent with a complete loss of function (Figure 1B). Development is delayed in both the *lonp-1(cmh24)* and *lonp-1(cmh25)* worms relative to wildtype worms (Figure 1C-E). To assess the impact of impaired LONP-1 on the mitochondrial network, we stained live worms with the fluorescent dye TMRE which accumulates within functional mitochondria in a manner requiring OXPHOS and an inner mitochondrial membrane potential. As expected, TMRE fluorescence was markedly reduced in *lonp-1(cmh24)* and *lonp-1(cmh25)* worms relative to wildtype worms, consistent with mitochondrial dysfunction (Figure 1F, G). *lonp-1(cmh24)* and *lonp-1(cmh25)* worms had reduced thrashing activity compared to wildtype worms indicating neuromuscular defects at the adult stage (Figure 1H).

**Figure 1:**
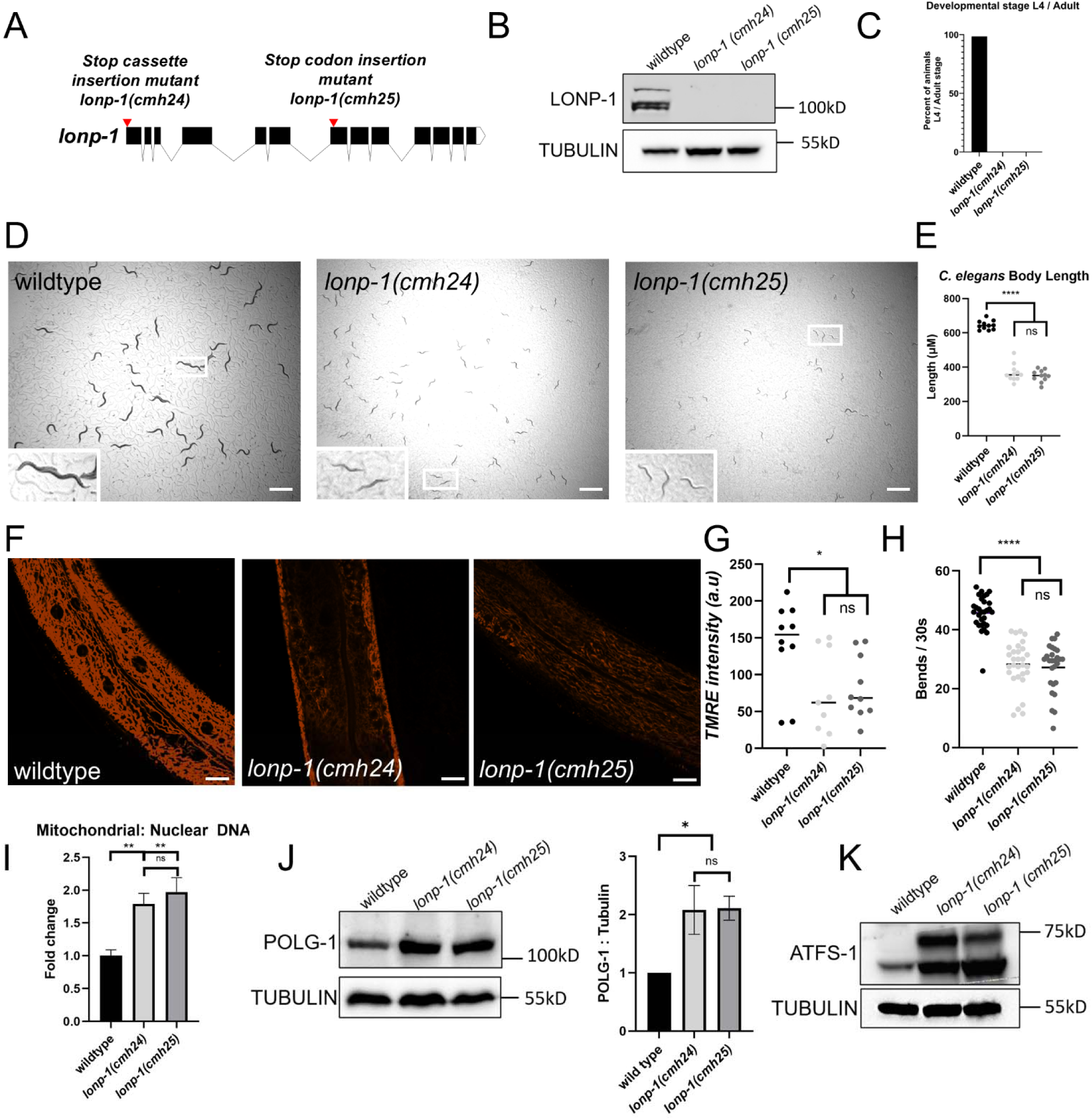
Mutants of *lonp-1* have severe mitochondrial dysfunction. **(A)** Schematic depicting location of stop codon mutations in *lonp-1*. **(B)** SDS-Page followed by western blot of wildtype, *lonp-1(cmh24)* and *lonp-1(cmh25)* worm lysates for LONP-1 and TUBULIN. **(C)** Percentage of wildtype, *lonp-1(cmh24)* and *lonp-1(cmh25)* worms that reach L4 or adult stage after 72 hours, n=200. **(D)** Representative brightfield images of wildtype, *lonp-1(cmh24)* and *lonp-1(cmh25)* worms after 72 hours at 11.3X magnification (scale bar, 1 mm). **(E)** Quantification of body length of wildtype, *lonp-1(cmh24)* and *lonp-1(cmh25)* worms after 72 hours, each point is a single animal, n=11 ****p<0.0001 (one-way ANOVA followed by Dunnett’s multiple comparison test). **(F)** Representative images of wildtype, *lonp-1(cmh24)* and *lonp-1(cmh25)* worms fluorescence after TMRE staining at 63X magnification. Scale bar 10 µm. **(G)** Quantification of fluorescence intensity of wildtype, *lonp-1(cmh24)* and *lonp-1(cmh25)* worms after TMRE staining, n=10 *p<0.05 (two-tailed Student’s *t*-test). a.u. Arbitrary units **(H)** Thrash assay of wildtype, *lonp-1(cmh24)* and *lonp-1(cmh25)* worms where each point is the number of body bends measured in an individual animal after 30 seconds, n=30 ****p<0.0001 (two-way ANOVA with post-hoc Sidak’s test). **(I)** Quantification of mtDNA by qPCR of wildtype, *lonp-1(cmh24)* and *lonp-1(cmh25)* worms, n=3 biological replicates of 25 worms. Error bars mean ± SD (two-tailed Student’s *t*-test) **p<0.005, NS, not significant. **(J)** Left: SDS-Page followed by western blot of wildtype, *lonp-1(cmh24)* and *lonp-1(cmh25)* worm lysates for POLG-1 and TUBULIN. Right: Quantification of western blot of POLG-1, n=3 biological replicates. Error bars mean ± SD (two-tailed Student’s *t*-test) *p<0.05, NS, not significant. **(L)** SDS-Page followed by western blot of *lonp-1* mutants for ATFS-1 and TUBULIN. All depicted experiments had three biological replicates with similar results.

mtDNA replication is tightly regulated, and mtDNA copy number is generally maintained in proportion to mitochondrial mass^20^. Interestingly, both *lonp-1(cmh24)* and *lonp-1(cmh25)* strains had a significant increase in mtDNA content compared to wildtype worms (Figure 1I). As POLG-1 is the sole polymerase required to replicate mtDNA^21^, we examined POLG-1 abundance in *lonp-1(cmh24)* and *lonp-1(cmh25)* worms compared to wildtype worms. Impressively, POLG-1 was increased 2-fold in the *lonp-1* mutants (Figure 1J). The increase in mtDNA copy number and POLG-1 abundance is consistent with previous work that suggests mtDNA replication is increased when *lonp-1* is impaired^22^. In *C. elegans,* mtDNA replication is increased when the bZIP protein ATFS-1 accumulates within mitochondria, which we observed in both *lonp-1(cmh24)* and *lonp-1(cmh25)* worms(Figure 1K)^23^. ATFS-1 regulates the UPR^mt^, which is a nuclear transcriptional program triggered by mitochondrial dysfunction^24^. Inhibition of *atfs-1* expression via RNAi in *lonp-1(cmh24)* and *lonp-1(cmh25)* worms caused an extended developmental delay, suggesting that *atfs-1* is protective during *lonp-1* inactivation (Supplemental Figure 1A). The increased ATFS-1 abundance and mtDNA copy number observed in the *lonp-1* mutants supports previous findings that ATFS-1 is a LONP-1 substrate that promotes mtDNA replication^22^.

Transcripts encoding mitochondrial proteases *spg-7, clpp-1, ppgn-1, and ymel-1* and mitochondrial chaperones *hsp-6* and *hsp-60* are reduced in *atfs-1(null)* worms, but elevated in *atfs-1(et18)*, a mutant strain in which ATFS-1 is constitutively localized to the nucleus (Supplementary Figure 1B)^25^. Via RNA-seq, we found that the same transcripts were increased in the *lonp-1(cmh24)* worms (Supplemental Figure 1C). Notably, crossing *atfs-1(gk3094)* loss of function mutants with *lonp-1* mutant alleles fails to produce viable homozygous offspring, indicating synthetic lethality when both are disrupted^26^. Consistent with previous studies, we conclude that ATFS-1 promotes survival and development during mitochondrial dysfunction caused by inhibition of LONP-1 by increasing mtDNA replication and by upregulating transcription of genes that support mitochondrial proteostasis and biogenesis.

### Mitochondrial ribosome mutations suppress *lonp-1* mutant phenotypes

To gain insight into the essential function of LONP-1, we performed a forward genetic screen to identify mutations that rescue the developmental delay caused by impaired *lonp-1* expression. *lonp-1(cmh25)* worms were mutagenized with ethyl methanesulfonate (EMS) and the F2 generation was screened for isolates that developed into adult worms in less than 100 hours (Figure 2A, B). Three isogenic suppressor strains were identified, outcrossed, and subjected to whole-genome sequencing. After sequencing, we identified point mutations in two genes. Two of the suppressor mutations were in the previously uncharacterized gene *Y54G9A.7* and the third suppressor was in *mrps-15*, which encodes a component of the small subunit of the mitochondrial ribosome (Figure 2C). *Y54G9A.7* contains a predicted mitochondrial targeting sequence and has been identified in proteomic datasets from *C. elegans* mitochondria (Supplemental Figure 2A)^27,28^. Phylogenetic analysis revealed that *Y54G9A.7* shares homology with the mammalian mitochondrial ribosomal subunit protein MRPS38, suggesting that *Y54G9A.7* encodes the *C. elegans* ortholog *mrps-38*^29^.

**Figure 2:**
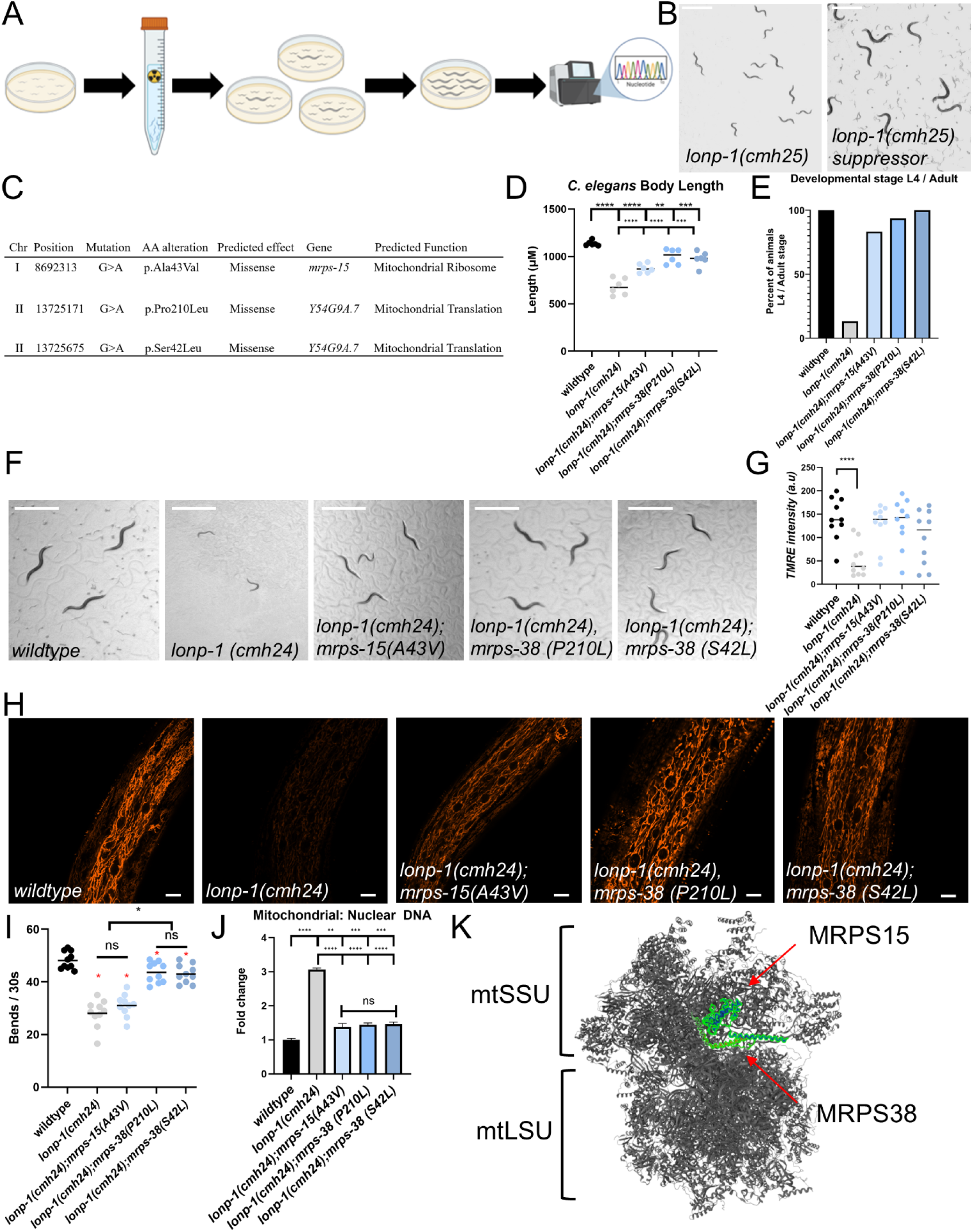
Mitochondrial ribosome mutations suppress *lonp-1* mutant phenotypes. **(A)** Schematic depicting mutagenesis screen for suppressors of *lonp-1(cmh25)* mutant growth phenotype. **(B)** Representative brightfield image of *lonp-1* suppressor mutant from screen after 72 hours at 11.3X magnification (scale bar, 1 mm). **(C)** Table of mutations identified from whole genome sequencing of *lonp-1* suppressor mutants. **(D)** Quantification of body length of wildtype, *lonp-1(cmh24)* and *lonp-1(cmh24)* with suppressor mutations after 72 hours, each point is a single animal, n=6. ****p<0.0001, ***p<0.001, **p<0.01 (one-way ANOVA followed by Dunnett’s multiple comparison test **(E)** Percentage of wildtype, *lonp-1(cmh24)* and *lonp-1(cmh24)* with suppressor mutations that reach L4 or adult stage after 72 hours, n=200. **(F)** Representative brightfield images of wildtype, *lonp-1(cmh24)* and *lonp-1(cmh24)* with suppressor mutations after 72 hours at 11.3X magnification (scale bar, 1 mm). **(G)** Quantification of fluorescence intensity of wildtype, *lonp-1(cmh24)* and *lonp-1(cmh24)* with suppressor mutations after TMRE staining, n=10. ****p<0.0001 (two-tailed Student’s *t*-test). a.u. Arbitrary units **(H)** Representative images of fluorescence of wildtype, *lonp-1(cmh24)* and *lonp-1(cmh24)* with suppressor mutations after TMRE staining at 63X magnification. Scale bar 10 µm. **(I)** Thrash assay of wildtype, *lonp-1(cmh24)* and *lonp-1(cmh24)* with suppressor mutations where each point is the number of body bends measured in an individual animal after 30 seconds, n=10 *p<0.01 (two-way ANOVA with post-hoc Sidak’s test). Red asterisks indicate significantly different than wildtype with p<0.01 **(J)** Quantification of mtDNA by qPCR of wildtype, *lonp-1(cmh24)* and *lonp-1(cmh24)* with suppressor mutations, n=3 biological replicates of 25 worms. Error bars mean ± SD (two-tailed Student’s *t*-test) ****p<0.0001, ***p<0.001, **p<0.01 NS, not significant. **(K)** Depiction of human mitochondrial ribosome small subunit proteins MRPS15 and MRPS38 adapted from PDB 6ZM5.^79^ All depicted experiments had three biological replicates with similar results.

CRISPR-cas9 genome editing was used to generate three novel *C. elegans* strains which harbor the suppressor mutations *mrps-38(P210L)*, *mrps-38(S42L)* and *mrps-15(A43V). mrps-38(P210L)*, *mrps-38(S42L)* and *mrps-15(A43V)* worms have comparable developmental rate, body size, TMRE staining, and mtDNA content to wildtype worms (Supplemental Figure 2B-D). However, the *mrps-38(P210L)* and *mrps-38(S42L)* worms exhibit a modest reduction in thrashing (Supplemental Figure 2E). The absence of overt phenotypes in *mrps-38(P210L)*, *mrps-38(S42L)* and *mrps-15(A43V)* worms suggests that these mutations exert context-specific effects related to defective mitochondrial proteostasis.

To validate the suppressor mutations identified in the EMS screen of *lonp-1(cmh25),* the *mrps-38* and *mrps-15* point mutations were introduced into a second strain, *lonp-1(cmh24)*. The resulting double mutants *lonp-1(cmh24);mrps-38(P210L)*, *lonp-1(cmh24);mrps-38(S42L)*, and *lonp-1(cmh24);mrps-15(A43V)* all developed more quickly than *lonp-1(cmh24)* worms, recapitulating the results from the mutagenesis screen (Figure 2D–F). We determined transcripts for developmental genes are reduced in *lonp-1(cmh24)* worms compared to wildtype worms and increased in *lonp-1(cmh24);mrps-38(P210)* worms compared to *lonp-1(cmh24)* worms from RNA-seq experiments (Supplemental Figure 2F-H, Supplemental Table 1). Developmental genes are not enriched in RNA-seq analysis of *mrps-38(P210)* worms compared to wildtype worms, suggesting that the *mrps-38(P210)* mutation does not restore growth by directly altering transcription of developmental genes (Supplemental Figure 2I, Supplemental Table 1).

Importantly, TMRE fluorescence was increased to wildtype levels in *lonp-1(cmh24)*;*mrps-38(P210L)*, *lonp-1(cmh24);mrps-38(S42L)* and *lonp-1(cmh24);mrps-15(A43V)* worms, suggesting that mitochondrial function was restored (Figure 2G, H). Furthermore, the elevated mtDNA copy number observed in *lonp-1(cmh24)* worms was significantly reduced in *lonp-1(cmh24)*;*mrps-38(P210L)*, *lonp-1(cmh24);mrps-38(S42L)* and *lonp-1(cmh24);mrps-15(A43V)* worms (Figure 2I). Together, the increased mitochondrial membrane potential and reduced mtDNA content are consistent with improved mitochondrial homeostasis. Furthermore, *lonp-1(cmh24)*;*mrps-38(P210L)*, and *lonp-1(cmh24);mrps-38(S42L)* adult worms have increased thrashing relative to *lonp-1(cmh24)* worms. However, thrashing remained low in the *lonp-1(cmh24);mrps-15(A43V)* worms, suggesting differences in the capacity of each mutation to restore organismal health in the absence of LONP-1 (Figure 2J).

We analyzed LONP-1 protein levels by western blot to test whether the suppressors act by restoring LONP-1 protein and determined LONP-1 abundance was unchanged in *lonp-1(cmh24)*;*mrps-38(P210L)*, *lonp-1(cmh24);mrps-38(S42L)* and *lonp-1(cmh24);mrps-15(A43V)* worms compared to *lonp-1(cmh24)* worms (Supplemental Figure 2J). Unlike the *mrps-38* point mutations, inhibition of *mrps-38* expression by RNAi did not result in faster development in *lonp-1* mutant worms (Supplemental figure 2K). Therefore, we suggest that the *mrps-38* mutations from the screen either confer a mild loss of function or alter MRPS-38 activity.

Interestingly, cryo-EM structural analysis of the human mitochondrial ribosome determined that MRPS-15 and MRPS-38 homologs are adjacent within the small subunit of the mitochondrial ribosome (Figure 2K)^30^. MRPS-15 and MRPS-38 are likely integrated in the *C. elegans* mitoribosomal small subunit in proximity with a shared or similar function. The identification of three unique mitochondrial ribosome mutations that can suppress *lonp-1* mutant phenotypes suggests a role for LONP-1 in regulation of translation within mitochondria.

### Inhibition of LONP-1 increases mitochondrial ribosome protein abundance and activity

The mitochondrial ribosome mutations identified in the suppressor screen of *lonp-1(cmh25)* worms led us to investigate mtDNA-encoded protein synthesis. We performed an *in organello* translation assay with mitochondria isolated from wildtype, *lonp-1(cmh24), lonp-1(cmh24)*;*mrps-38(P210L)*, or *mrps-38(P210L)* worms. Mitochondria purified from each strain were incubated with S^35^-labeled methionine and cysteine which is incorporated into newly synthesized proteins. Radiolabeled products were separated by SDS-PAGE and visualized by phosphor-imaging. Unexpectedly, *lonp-1(cmh24)* mitochondria exhibited increased radiolabeled protein compared to wildtype mitochondria, indicating increased mitochondrial translation (Figure 3A). In contrast, *lonp-1(cmh24);mrps-38(P210L)* mitochondria had reduced radiolabeling relative to *lonp-1(cmh24)* mitochondria, suggesting that the suppressor mutation decreases protein synthesis within mitochondria (Figure 3A).

**Figure 3:**
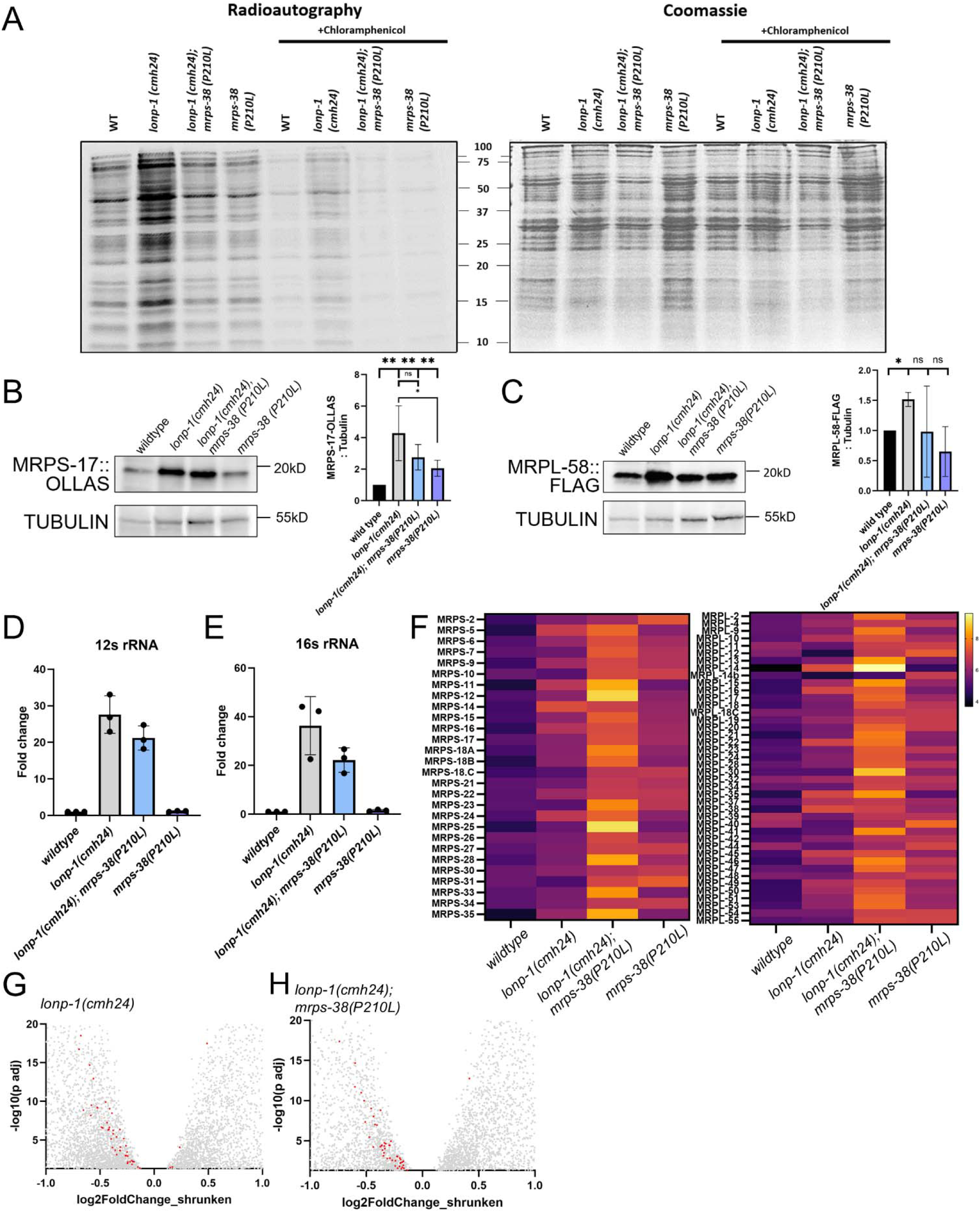
LONP-1 deficiency increases mitochondrial ribosome protein abundance and activity. **(A)** ^35^S-methionine and ^35^S-cysteine metabolic labeling of mitochondria isolated from wildtype, *lonp-1(cmh24)*, *lonp-1(cmh24);mrps-38(P210L)* and *mrps-38(P210L)* worms for 60 min. in the presence of emetine and cycloheximide (100 ug/mL) followed by SDS-Page and autoradiography, samples were treated with 200 μM chloramphenicol (mitochondrial ribosome inhibitor) as a control. Coomassie staining was used as a loading control. **(B)** Left: SDS-Page followed by western blot of *mrps-17(OLLAS)*, *lonp-1(cmh24);mrps-17(OLLAS)*, *mrps-17(OLLAS);mrps-38(P210L);lonp-1(cmh24)* and *mrps-17(OLLAS);mrps-38(P210L)* worm lysates for OLLAS and TUBULIN. Right: Quantification of western blot of OLLAS, n=3 biological replicates. Error bars mean ± SD (two-tailed Student’s *t*-test) *p<0.05, **p<0.005 NS, not significant **(C)** Left: SDS-Page followed by western blot of *mrpl-58(FLAG)*, *lonp-1(cmh24);mrpl-58(FLAG)*, *mrpl-58(FLAG);mrps-38(P210L);lonp-1(cmh24)* and *mrpl-58(FLAG);mrps-38(P210L)* worm lysates for FLAG and TUBULIN. Right: Quantification of western blot of OLLAS, n=3 biological replicates. Error bars mean ± SD (two-tailed Student’s *t*-test) *p<0.05, NS, not significant **(D)** Fold change in transcripts from RNA-seq of wildtype, *lonp-1(cmh24)*, *lonp-1(cmh24);mrps-38(P210L)* and *mrps-38(P210L)* worms for MTCE.7 (12s ribosomal RNA) **(E)** Fold change in transcripts from RNA-seq of wildtype, *lonp-1(cmh24)*, *lonp-1(cmh24);mrps-38(P210L)* and *mrps-38(P210L)* worms for MTCE.33 (16s ribosomal RNA) **(F)** Label-free quantification (LFQ) intensity heat maps of large and small mitochondrial ribosome proteins from mitochondrial isolated from wildtype, *lonp-1(cmh24)*, *lonp-1(cmh24);mrps-38(P210L)* and *mrps-38(P210L)* worms. n=4 MS experiments **(G)** Volcano plot of transcripts measured by RNA-seq from *lonp-1(cmh24)* worms, mitochondrial ribosome proteins are colored red n=3 (H) Volcano plot of transcripts measured by RNA-seq from *lonp-1(cmh24);mrps-38(P210L)* worms, mitochondrial ribosome proteins are colored red n=3. All depicted experiments had three biological replicates with similar results.

We hypothesized that the increased mitochondrial protein synthesis in *lonp-1(cmh24)* was caused by an increase in the abundance of mitochondrial ribosomes. In order to determine mitochondrial ribosome protein abundance, we used CRISPR-Cas9 genome editing to generate a *C. elegans* strain with epitope tags on two mitochondrial ribosome subunits: a FLAG tag at the C-terminus of the large subunit protein MRPL-49, and an OLLAS tag at the C-terminus of the small subunit protein MRPS-17. Whole animal lysate was analyzed by western blot which showed increased levels of both MRPL-49::FLAG and MRPS-17::OLLAS in *lonp-1(cmh24)* worms compared to wildtype worms (Figure 3B,C). We next examined mtDNA-encoded ribosomal RNA (mt-rRNA) which is a core mitochondrial ribosome component. We analyzed transcripts by RNA-seq and determined mt-rRNAs are also elevated in *lonp-1(cm24)* worms compared to wildtype worms, consistent with the observed increase in mitochondrial ribosomal proteins (Figure 3D,E).

Metazoan mitochondrial ribosome proteins are encoded and transcribed in the nucleus, translated in the cytosol and imported into mitochondria where they are assembled into mitoribosomes or degraded^31^. We performed quantitative mass spectrometry on mitochondria isolated from wildtype, *lonp-1(cmh24), lonp-1(cmh24)*;*mrps-38(P210L),* and *mrps-38(P210L)* worms and determined that mitochondria isolated from *lonp-1(cmh24)* or *lonp-1(cmh24)*;*mrps-38(P210L*) mitochondria had elevated levels of many mitochondrial ribosome proteins (Figure 3F). Interestingly, from RNA-seq analysis we found that transcript levels of nearly all mitochondrial ribosome genes in *lonp-1(cmh24)* worms, as well as *lonp-1(cmh24)*;*mrps-38(P210L)* worms, are reduced compared to wildtype worms (Figure 3G,H).

The observation that *lonp-1(cmh24)* worms harbor increased mitochondrial ribosomal proteins while their corresponding mRNA transcripts are reduced suggests that there is a reduction in protein degradation, and not an upregulation of mitoribosomal genes. We compared data from yeast, mouse and human cell line models of LONP-1 deficiency with our mass spectrometry results from LONP-1 deficient strains *lonp-1(cmh24)* and *lonp-1(cmh24);mrps-38(P210L)* and generated a list of putative *C. elegans* LONP-1 substrates, which feature many mitochondrial ribosomal proteins (Supplemental Table 2)^32–34^. We therefore propose that undegraded mitochondrial ribosomal proteins accumulate in the absence of LONP-1. Further, we suggest that the increase in mtDNA-encoded protein synthesis in *lonp-1(cmh24)* worms is due to an increased number of mitoribosomes caused by increased biogenesis, resulting from the highly abundant mt-rRNA and undegraded mitochondrial ribosome protein components.

### mrps-38(P210L), mrps-38(S42L) and mrps-15(A43V) mutations restore impaired OXPHOS in lonp-1(cmh24) worms

We next examined whether mutations in *lonp-1*, *mrps-15* and *mrps-38* affected the abundance of mtDNA-encoded proteins. We first measured the abundance of the cytochrome c oxidase subunit 1 (MTCOI), which is encoded by mtDNA and synthesized on mitochondrial ribosomes as an early step in complex IV biogenesis^35^. Surprisingly, MTCOI protein is reduced in *lonp-1(cmh24)* worms compared to wildtype worms, despite the previously observed increase in mtDNA-encoded protein synthesis (Figure 4A, Supplemental Figure 4A,B). In contrast, MTCOI levels in *lonp-1(cmh24);mrps-38(P210L)*, *lonp-1(cmh24);mrps-38(S42L)*, and *lonp-1(cmh24);mrps-15(A43V)* worms are comparable to those observed in wildtype worms (Figure 4A, Supplemental Figure 4A,B). Notably, MTCOI abundance is unchanged in *mrps-15(A43V)*, *mrps-38(S42L)*, and *mrps-38(P210L)* worms, indicating that these mutations do not impact MTCOI under basal conditions (Figure 4A, Supplemental Figure 4A,B).

**Figure 4.**
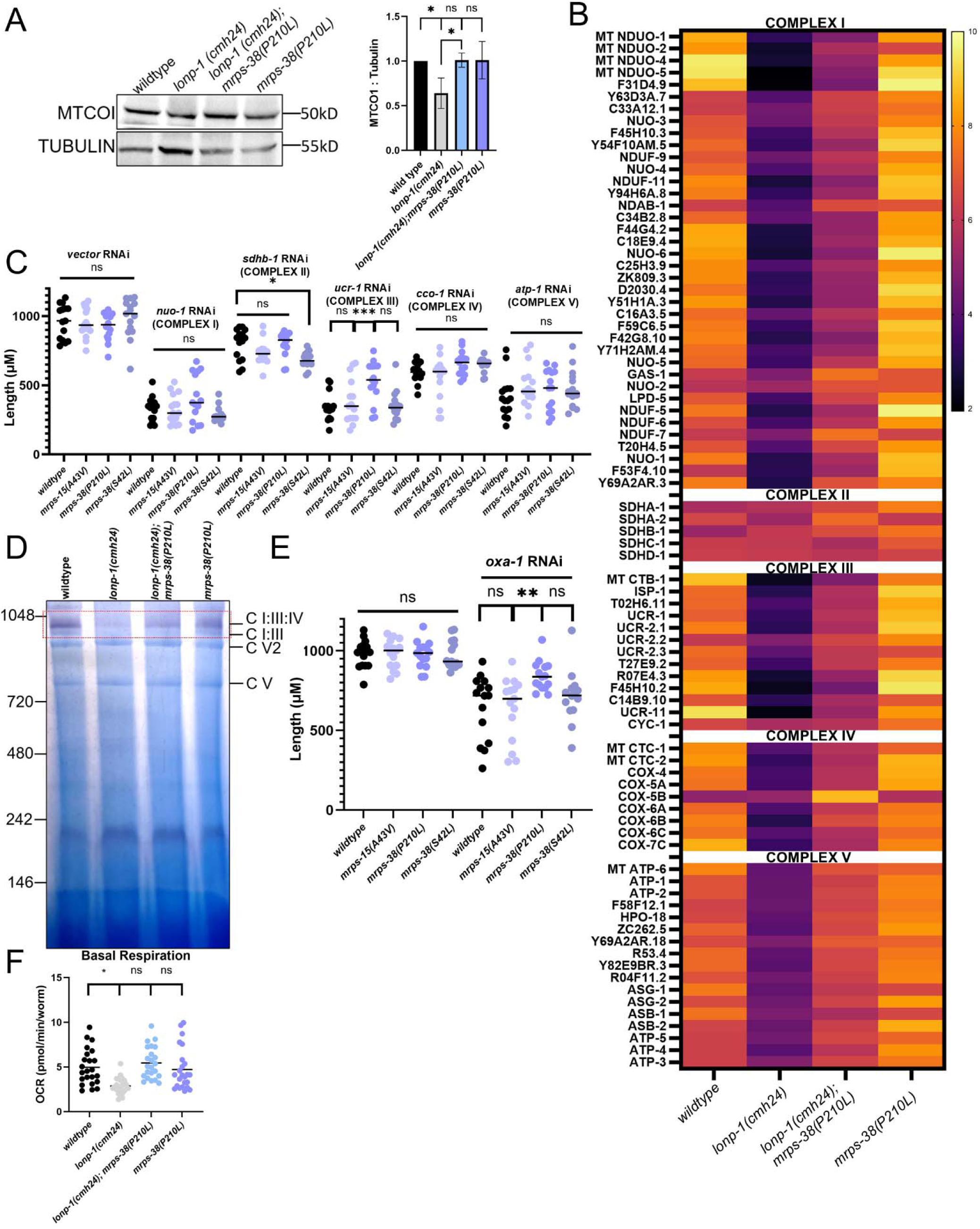
mrps-38(P210L), mrps-38(S42L) and mrps-15(A43V) restore impaired OXPHOS in lonp-1(cmh24) worms. **(A)** Left: SDS-Page followed by western blot of wildtype, *lonp-1(cmh24), mrps-38(P210L);lonp-1(cmh24)* and *mrps-38(P210L)* worm lysates for MT-COI and TUBULIN. Right: Quantification of western blot of MT-COI, n=3 biological replicates. Error bars mean ± SD (two-tailed Student’s *t*-test) *p<0.05, **p<0.005 NS, not significant **(B)** Label-free quantification (LFQ) intensity heat maps of OXPHOS proteins from mitochondrial isolated from wildtype, *lonp-1(cmh24)*, *lonp-1(cmh24);mrps-38(P210L)* and *mrps-38(P210L)* worms. n=4 MS experiments **(C)** Quantification of body length of wildtype, *mrps-15(A43V), mrps-38(P210L) and mrps-38(S42L)* worms after 72 hours of growth on RNAi, each point is a single animal, n=15 *p<0.05, ***p<0.001 (one-way ANOVA followed by Dunnett’s multiple comparison test) NS, not significant **(D)** BN-PAGE of mitochondria isolated from wildtype, *lonp-1(cmh24)*, *lonp-1(cmh24);mrps-38(P210L)* and *mrps-38(P210L)* worms followed by complex I in-gel activity assay **(E)** Quantification of body length of wildtype, *mrps-15(A43V), mrps-38(P210L) and mrps-38(S42L)* worms after 72 hours of growth on *oxa-1* RNAi, each point is a single animal, n=15 *p<0.05, ***p<0.001 (one-way ANOVA followed by Dunnett’s multiple comparison test) NS, not significant **(F)** Oxygen consumption rates (OCR) of wildtype, *lonp-1(cmh24)*, *lonp-1(cmh24);mrps-38(P210L)* and *mrps-38(P210L)* worms, each point represents 1 well of 10 animals, *p<0.05, NS, not significant, (two-tailed Student’s t-test). All depicted experiments had three biological replicates with similar results.

We performed GO analysis of quantitative mass spectrometry data from mitochondria isolated from wildtype, *lonp-1(cmh24), lonp-1(cmh24)*;*mrps-38(P210L),* and *mrps-38(P210L)* worms for all putative mitochondrial proteins, determined from previous *C. elegans* experiments and MitoCarta homology^27,36,37^ (Supplemental Table 3). We found electron transport chain components were the most significantly changed GO category between *lonp-1(cmh24*) and wildtype worms, as well as in the comparison of *lonp-1(cmh24)* and *lonp-1(cmh24)*;*mrps-38(P210L)* (Supplemental Figure 4C-E, Supplemental Table 4). We then examined mass spectrometry data for all *C. elegans* homologs of human OXPHOS proteins and found nearly all measurable OXPHOS proteins were reduced in mitochondria from *lonp-1(cmh24)* worms compared to mitochondria from wildtype worms (Figure 4D, Supplemental Table 5). Complex II is the only OXPHOS complex that is entirely encoded by nuclear DNA^2^. Notably, the proteins that comprise complex II appear unchanged, suggesting that only OXPHOS complexes with mtDNA-encoded components are reduced by LONP-1 deficiency (Figure 4D). Importantly, *lonp-1(cmh24);mrps-38(P210L)* mitochondria have increased OXPHOS proteins compared to *lonp-1(cmh24)* mitochondria (Figure 4D). Combined, these results suggest that inactivation of *lonp-1* causes a reduction in OXPHOS proteins which is prevented or rescued by mutations in *mrps-38* or *mrps-15*.

To determine if the mitochondrial ribosome mutations could restore OXPHOS defects, we inhibited subunits from each of the five respiratory chain complexes: *nuo-1* (complex I), *sdha-1* (complex II), *ucr-1* (complex III), *cco-1* (complex IV), and *atp-1* (complex V) via RNAi. Knockdown of these OXPHOS genes reduced development of wildtype worms, *mrps-38(P210L)*, *mrps-38(S42L)*, and *mrps-15(A43V)* mutants equally with the exception of *mrps-38(P210L)* treated with *ucr-1* RNAi (Figure 4E, Supplemental Figure 4F). The developmental delay caused by RNAi inhibition of OXPHOS suggests that the suppressor mutations cannot directly restore reduced OXPHOS proteins. Therefore, we propose that the *mrps-38(P210L)*, *mrps-38(S42L)*, and *mrps-15(A43V)* mutants suppress the developmental delay caused by LONP-1 deficiency by enhancing assembly of OXPHOS complexes, which may prevent the reduction of OXPHOS protein levels observed in *lonp-1* mutants.

Previous findings in yeast and mammalian cells established a role for LONP-1 in promoting assembly of OXPHOS complexes, acting as a protein chaperone via the ATPase domain^12,13^. We hypothesized that the reduced OXPHOS protein abundance in *lonp-1(cmh24)* worms reflects a defect in OXPHOS complex assembly which results in rapid degradation of unassembled OXPHOS proteins^5^. To analyze the OXPHOS complexes, we performed blue native PAGE on digitonin-treated mitochondria isolated from wildtype, *lonp-1(cmh24), lonp-1(cmh24)*;*mrps-38(P210L)*, and *mrps-38(P210L)* worms. The gel was then incubated with nitroblue tetrazolium, which stains purple in the presence of active complex I. As expected, complex I activity was significantly reduced in *lonp-1(cmh24)* mitochondria relative to wildtype mitochondria, suggesting that complex I assembly is reduced when LONP-1 is impaired (Figure 4F). Interestingly, we observed a significant increase in complex I activity in *lonp-1(cmh24);mrps-38(P210L)* mitochondria compared to *lonp-1(cmh24)* mitochondria, suggesting that the *mrps-38(P210L)* mutation can prevent or recover defective OXPHOS assembly (Figure 4F). No significant difference in complex I was observed between wildtype and *mrps-38(P210L)* worms and GO analysis of mass spectrometry from *mrps-38(P210L)* mitochondria did not identify any OXPHOS-related categories, suggesting that the mutation does not increase complex I assembly in basal conditions (Figure 4F, Supplemental Figure 4G, Supplemental Table 4).

To assess whether the mitoribosome mutations promote OXPHOS complex assembly, we examined the sensitivity of *mrps-38(P210L)*, *mrps-38(S42L)*, and *mrps-15(A43V)* mutants to RNAi inhibition of *oxa-1*, a mitochondrial translocase localized within the mitochondrial inner membrane(IM)^38^. OXA-1 interacts with mitoribosomes to mediate co-translational protein folding and insertion of nascent proteins into the IM, thereby facilitating OXPHOS assembly^39,40^. RNAi knockdown of *oxa-1* caused impaired growth of wildtype worms, which was attenuated in *mrps-38(P210L)* worms, suggesting the suppressor mutation can partially rescue defects in OXPHOS assembly (Figure 4G, Supplemental Figure 4H).

To gain insight into how LONP-1 and MRPS-38 impact respiration, we measured oxygen consumption in wildtype, *lonp-1(cmh24), lonp-1(cmh24)*;*mrps-38(P210L)*, and *mrps-38(P210L)* worms. Basal respiration was significantly reduced in *lonp-1(cmh24)* worms relative to wildtype worms but was increased to wildtype levels in *lonp-1(cmh24);mrps-38(P210L)* worms (Figure 4H). The restoration of respiration in *lonp-1(cmh24)* worms by the *mrps-38(P210L)* mutation suggests that altering the mitochondrial ribosome can be beneficial when the electron transport chain is disrupted.

The reduction of OXPHOS protein abundance, complex I activity and respiration observed in *lonp-1(cmh24)* worms suggests that in the absence of LONP-1, OXPHOS is severely impaired, which likely explains the mitochondrial and organismal dysfunction observed in the *lonp-1* mutant worms. The restoration of OXPHOS assembly and activity by a suppressor mutation within the mitochondrial ribosome suggests that protein synthesis on mitoribosomes is dysregulated in the absence of LONP-1, and that reducing mitochondrial translation can enhance OXPHOS complex assembly during LONP-1 dysfunction.

### The *mrps-38(P210L)* point mutation reduces protein synthesis on mitochondrial ribosomes

*lonp-1(cmh24)*;*mrps-38(P210L)* worms do not exhibit the elevated protein synthesis observed in *lonp-1(cmh24)* mitochondria, suggesting that the *mrps-38(P210L*) mutation causes a reduction in mtDNA-encoded protein synthesis (Figure 3A). MRPL-49::FLAG and MRPS-17::OLLAS are both increased in *lonp-1(cmh24)*;*mrps-38(P210L)* worms compared to wildtype worms (Figure 3B,C). Mass spectrometry indicates increased accumulation of mitochondrial ribosome proteins in *lonp-1(cmh24);mrps-38(P210L)* mitochondria relative to wildtype mitochondria (Figure 3F). *lonp-1(cmh24);mrps-38(P210L)*worms also have increased transcript levels of mt-rRNAs relative to wildtype worms (Figure 3D,E). Taken together, the observation that *mrps-38(P210L);lonp-1(cmh24)* worms have wildtype levels of protein synthesis with elevated levels of mitochondrial ribosomal proteins and RNAs suggests that the *mrps-38(P210L)* mutation reduces mitochondrial translation. Furthermore, mitochondria from *mrps-38(P210L)* worms had increased mitoribosomal protein levels but had reduced levels of radiolabeled protein compared to wildtype mitochondria in the mitochondrial translation assay (Figure 3A,F). These findings suggest that the *mrps-38(P210L)* mutation does not reduce ribosome abundance but instead slows the rate of translation, leading to lower mtDNA-encoded protein synthesis despite elevated mitoribosomal protein levels.

The human homolog of MRPS-38 associates with proteins and RNA that form the small subunit of the mitochondrial ribosome early in the assembly process.^41^ Therefore, we sought to determine if the *mrps-38(P210L*) mutation altered mitochondrial ribosome assembly. We performed sucrose gradient fractionation on mitochondria isolated from wildtype, *lonp-1(cmh24)*, *lonp-1(cmh24);mrps-38(P210L)*, and *mrps-38(P210L)* worms expressing MRPL-49::FLAG and MRPS-17::OLLAS. Analysis of gradient fractions which corresponded with assembled mitoribosomes by western blot revealed comparable levels of MRPL-49::FLAG and MRPS-17::OLLAS in all strains (Figure 5A). Therefore, we conclude that the *mrps-38(P210L)* mutation does not alter mitochondrial ribosome assembly. We observed no protein in the light fractions from *lonp-1(cmh24)* mitochondria, which contrasts from wildtype, *lonp-1(cmh24);mrps-38(P210L)*, and *mrps-38(P210L)* mitochondria, which suggests that there is very little unassembled MRPL-49 and MRPS-17 (Figure 5A).

**Figure 5.**
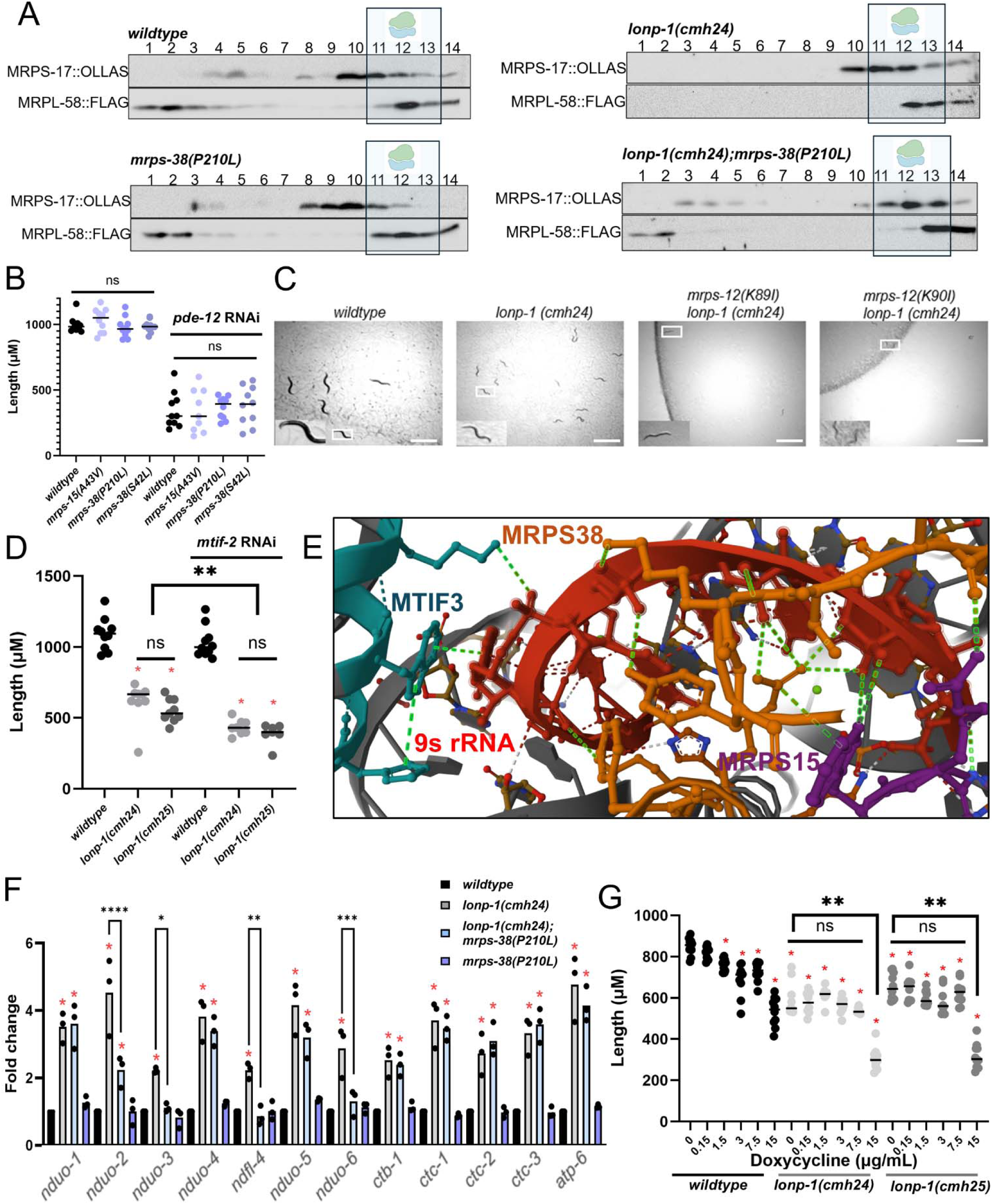
*mrps-38(P210L)* point mutation reduces protein synthesis by the mitochondrial ribosome. **(A**) Anti-FLAG and Anti-OLLAS western blots of sucrose density gradient fractions from mitochondria isolated from of *mrps-17(OLLAS);mrpl-58(FLAG)*, *mrps-17(OLLAS);lonp-1(cmh24);mrpl-58(FLAG)*, *mrps-17(OLLAS);mrpl-58(FLAG);mrps-38(P210L);lonp-1(cmh24)* and *mrps-17(OLLAS);mrpl-58(FLAG);mrps-38(P210L)*worms. Gradient fractions 11-13 represent monosomes. **(B)** Quantification of body length of wildtype, *mrps-15(A43V), mrps-38(P210L) and mrps-38(S42L)* worms after 72 hours of growth on *pde-12* RNAi, each point is a single animal, n=10 (one-way ANOVA followed by Dunnett’s multiple comparison test) NS, not significant **(C)** Representative brightfield images of wildtype, *lonp-1(cmh24), lonp-1(cmh24);mrps-12(K89I)* and *lonp-1(cmh24);mrps-12(K90I)* worms after 72 hours at 11.3X magnification (scale bar, 1 mm) **(D)** Quantification of body length of wildtype, *lonp-1(cmh24)* and *lonp-1(cmh25)* worms after 72 hours growth on *mtif-2* RNAi, each point is a single animal, n=10 **p<0.001 (one-way ANOVA followed by Dunnett’s multiple comparison test **(E)** Cryo-EM structure of *Trypanosoma brucei* small mitoribosomal subunit in complex with mt-IF-3 PDB: 6HIW^80^ MTIF3 is colored green, 9s rRNA is colored red, MRPS38 is colored orange and MRPS15 is colored purple. Putative interactions are highlighted in green. **(F)** Fold change in transcripts from RNA-seq of wildtype, *lonp-1(cmh24)*, *lonp-1(cmh24);mrps-38(P210L)* and *mrps-38(P210L)* worms for mtDNA-encoded genes. Red asterisks denote significantly different from wildtype *p<0.05, **p<0.01 ***p<0.0005, ****p<0.0001 (one-way ANOVA followed by Dunnett’s multiple comparison test) **(G)** Quantification of body length of wildtype, *lonp-1(cmh24)* and *lonp-1(cmh25)* worms after 72 hours growth on doxycycline doses 0, 0.15, 1.5, 3, 7.5, and 15 µg/mL, each point is a single animal, n=10. Red asterisks denote significantly different from wildtype *p<0.05, **p<0.001 (one-way ANOVA followed by Sidak’s multiple comparison test). All depicted experiments had three biological replicates with similar results.

Increased mitochondrial ribosome activity could lead to an increase in stalling, potentially reducing functional OXPHOS protein synthesis despite increased mitochondrial translation. We tested the susceptibility of *mrps-38(P210L)*, *mrps-38(S42L)*, and *mrps-15(A43V)* worms to RNAi targeting the gene homologous to human PDE12, which regulates polyadenylation of mitochondrial RNAs and causes mitochondrial ribosome stalling in its absence^42^ (Figure 5B). We observed no difference in response to treatments that induce ribosome stalling in *mrps-38(P210L)*, *mrps-38(S42L)*, and *mrps-15(A43V)* worms compared to wildtype worms, suggesting that the suppressor mutations do not increase or decrease mitochondrial ribosome stalling.

The fidelity of translation within mitochondria is dependent on interactions that pair charged tRNA and cognate amino acid within the mitochondrial ribosome^43^. It has been observed in bacteria, yeast and mice that mutations in the mitochondrial ribosome proteins homologous to *C. elegans* MRPS-12 can increase or decrease mitochondrial translation fidelity^44^. *C. elegans* strains with homologous mutations to those that cause hyper-accurate (*mrps-12(K89T))* and error-prone *(mrps-12(K90I))* mitochondrial translation were generated previously^45^. We crossed worms with these point mutations in *mrps-12* into the *lonp-1(cmh24)* background and found the *lonp-1(cmh24);mrps-12* mutant worms develop slowly (Figure 5C). The *lonp-1(cmh24);mrps-12* mutant worms do not phenocopy *lonp-1(cmh24);mrps-38(P210L)* suggesting that the *mrps-38(P210L)* mutation does not significantly impact translation fidelity within mitochondria.

Mitochondrial ribosomes initiate translation by forming a pre-initiation complex that includes the small subunit of the mitochondrial ribosome, tRNA methionine, mtDNA-encoded mRNAs (mt-mRNA) and the initiation factors MTIF2 and MTIF3^46^. Inhibition of MTIF2 by RNAi in *lonp-1(cmh24)* worms exacerbates the developmental delay, suggesting that altered mitochondrial ribosome translation initiation is not the mechanism that suppresses *lonp-1(cmh24)* mitochondrial dysfunction (Figure 5D). We speculate that the elevated mtDNA-encoded protein synthesis in *lonp-1(cmh24)* worms supports growth, perhaps by enabling a minimal level of OXPHOS assembly from the enriched mtDNA-encoded protein synthesis. The C-terminal region of *C. elegans* MRPS-38 where the P210L mutation is found does not appear to be conserved in yeast or mammals however it occurs in *Trypanosoma brucei* and *Trypanosoma cruzi* (Supplemental Figure 5A). Cryo-EM structures of *Trypanosoma brucei* and *Trypanosoma cruzi* mitoribosomes show a shared region of interaction between mitochondrial ribosomal RNA, the C-terminal of the MRPS-38 homolog MRPS38 and MTIF3 (Figure 5E, Supplemental Figure 5B,C). CRISPR knock out of MTIF3 causes a significant increase in mtDNA-encoded protein synthesis, suggesting that MTIF3 negatively regulates mitochondrial translation initiation^47^. Although MTIF3 has not been identified in *C. elegans*, its strong evolutionary conservation suggests that a functional homolog may exist.^48^ Therefore, we cannot rule out the possibility that MRPS-38(P210L) suppresses mitochondrial dysfunction via interaction with an MTIF3-like factor.

Mitochondrial encoded mRNA transcripts are elevated in *lonp-1(cmh24)* worms compared to wildtype worms (Figure 5F). mt-mRNA transcripts are similarly increased in *lonp-1(cmh24);mrps-38(P210L)* worms compared to wildtype worms, with the exception of transcripts encoding protein components of complex I identified as *nduo-2*, *nduo-3*, *ndfl-4* and *nduo-6* (Figure 5F). We observed a uniform decrease in protein bands in the *in organello* translation assay, which suggests that the reduction in translation is not specific to *nduo-2*, *nduo-3*, *ndfl-4* and *nduo-6* (Figure 3A). Further, NDUO-2 protein is not reduced in *lonp-1(cmh24);mrps-38(P210L)* mitochondria compared to *lonp-1(cmh24*) mitochondria (Figure 4D). Taken together, the results suggest that the *mrps-38(P210L)* mutation does not reduce mitochondrial translation by reducing transcription of mt-mRNA. It should be noted that *nduo-3*, *ndfl-4* and *nduo-6* were not identified in the mass spectrometry experiment (Figure 4D). A recent model of complex I assembly proposes that formation of a module consisting of ND2, ND3, ND4L and ND6 (homologous proteins to *C. elegans nduo-2*, *nduo-3*, *ndfl-4* and *nduo-6)* is the initial step in the biogenesis of complex I^49^. We propose that the reduction in *nduo-2*, *nduo-3*, *ndfl-4* and *nduo-6* transcripts in *mrps-38(P210L);lonp-1(cmh24*) mitochondria compared to *lonp-1(cmh24*) mitochondria is due to post-translational mRNA decay following successful assembly of the complex I module.

Two competing scenarios could explain how the MRPS-38(P210L) mutation reduces translation within mitochondria of *lonp-1(cmh24)* worms. One possibility is that the ribosome population is heterogeneous, with a subset of mitochondrial ribosomes functioning normally while a subset of assembled mitochondrial ribosomes are inactive. Alternatively, all mitochondrial ribosomes may remain functional but operate at a reduced rate. To distinguish between these models, we treated *lonp-1* mutants with doxycycline, an inhibitor of mitochondrial translation. Because chemical inhibitors act by directly binding ribosomes, a low dose of ribosome inhibitor will result in a subset of functional ribosomes that remain unbound by the inhibitor. However, we did not identify a concentration of ribosome inhibition that restored development in the *lonp-1* mutants to that of wildtype worms, suggesting that partial inhibition does not mimic the MRPS-38(P210L) effect (Figure 5G). Thus, we favor the interpretation that the mutation slows mitochondrial translation globally rather than in only a subset of ribosomes.

### C. elegans models of mitochondrial disease are improved by mitoribosome mutations

Cerebral, ocular, dental, auricular and skeletal syndrome (CODAS) is a rare disease caused by mutations in LONP-1^18^. One homozygous mutation that has been identified as a CODAS allele is LONP1(R721G)^50^. We generated a homologous *lonp-1(R711G)* mutation in *C. elegans* using CRISPR-cas9 genome editing (Figure 6A). We then crossed the *lonp-1(R711G)* worms with *hsp-6_pr_::GFP* worms to monitor UPR^mt^ activation as an indicator of mitochondrial dysfunction. As expected, *hsp-6_pr_::GFP* expression was increased in *lonp-1(R711G)*;*hsp-6_pr_::GFP* worms relative to *hsp-6_pr_::GFP* worms, indicating activation of the UPR^mt^ and mitochondrial dysfunction (Figure 6B). *lonp-1(R711G)* adult worms had reduced thrashing compared to wildtype adult worms, suggesting a defect in muscular function or coordination (Figure 6C). We then crossed *lonp-1(R711G)* worms with either *mrps-38(P210L)* worms or *mrps-38(S42L)* worms (we were unable to cross with *mrps-15(A43V)* worms due to their shared chromosomal region). We found that *lonp-1(R711G)*;*mrps-38(P210L)* and *lonp-1(R711G)*;*mrps-38(S42L*) worms had increased levels of thrashing compared to *lonp-1(R711G)* worms, which was similar to the thrashing rate of wildtype worms (Figure 6C). *lonp-1(R711G)* worms also exhibited reduced egg laying compared to wild type worms, which was increased in *lonp-1(R711G)*;*mrps-38(P210L)* and *lonp-1(R711G)*;*mrps-38(S42L*) worms (Figure 6D).

**Figure 6.**
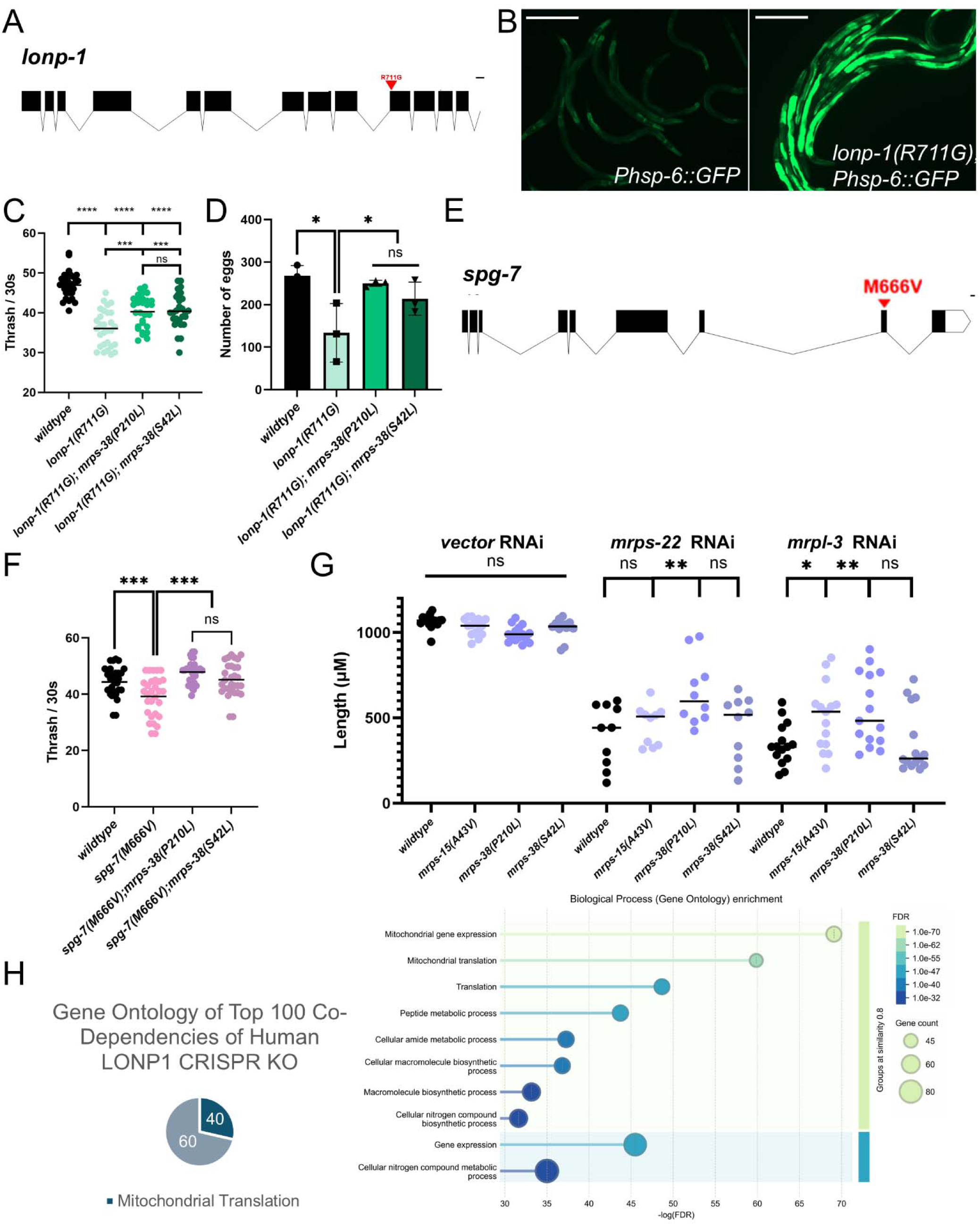
*mrps-38(P210L) and mrps-38(S42L)* ameliorate disease models of CODAS and SCA28. **(A)** Schematic depicting location of CODAS-linked homologous mutation in *lonp-1* **(B)** Representative image of *Phsp-6::GFP* and *lonp-1(R711G);Phsp-6::GFP* worms’ GFP fluorescence after 48 hours (scale bar, 0.5mm) **(C)** Thrash assay of wildtype, *lonp-1(R711G)*, *lonp-1(R711G);mrps-38(P210L), and lonp-1(R711G);(mrps-38(S42L)* worms where each point is the number of body bends measured in an individual animal after 30 seconds, n=30 ***p<0.0005 ****p<0.0001 NS, not significant (two-way ANOVA with post-hoc Sidak’s test). **(D)** Number of eggs laid by wildtype, *lonp-1(R711G)*, *lonp-1(R711G);mrps-38(P210L), and lonp-1(R711G);(mrps-38(S42L)* worms *p<0.05, ns, not significant (one-way ANOVA followed by Tukey’s multiple comparison test) **(E)** Schematic depicting location of SCA28-linked homologous mutation in *spg-7* **(F)** Thrash assay of wildtype, *spg-7(M666V)*, *spg-7(M666V);mrps-38(P210L), spg-7(M666V);(mrps-38(S42L)* worms where each point is the number of body bends measured in an individual animal after 30 seconds, n=30 ***p<0.0005 NS, not significant (two-way ANOVA with post-hoc Sidak’s test). **(G)** Quantification of body length of wildtype, *mrps-15(A43V), mrps-38(P210L) and mrps-38(S42L)* worms after 72 hours of growth on *mrps-22* or *mrpl-3* RNAi, each point is a single animal, n=10 *p<0.05 **p<0.005 NS, not significant (one-way ANOVA followed by Dunnett’s multiple comparison test) **(H)** GO analysis of the Top 100 Co-dependencies of human LONP1 CRISPR KO from DepMap^56^. All depicted experiments had three biological replicates with similar results.

We then investigated whether the mitoribosome suppressor mutations could correct other models of disease related to mitochondrial proteostasis. Spinocerebellar ataxia 28 (SCA28) is a disease characterized by slow progressive ataxia and cerebellum shrinking which occurs later in life^51^. SCA28 is caused by mutations in a gene encoding the inner membrane-localized mitochondrial protease AFG3L2 which degrades misfolded proteins and promotes OXPHOS assembly^52^. We generated a *C. elegans* strain harboring a homologous mutation to that found in SCA28 patients, identified as *spg-7(M666V)* (Figure 6E). *spg-7(M666V)* adult worms exhibited reduced thrashing compared to wildtype adult worms, suggesting defective neuromuscular function (Figure 6E). We crossed *spg-7(M666V)* worms with *mrps-38(P210L)* or *mrps-38(S42L)* worms and found that the thrashing rate was increased in *spg-7(M666V)*;*mrps-38(P210L)* and *spg-7(M666V*);*mrps-38(S42L)* worms (Figure 6F). We were unable to cross with *mrps-15(A43V)* worms due to their shared chromosomal region.

Numerous diseases are caused by mutations in mitochondrial ribosome genes, which can impact OXPHOS protein production and assembly^53^. We examined the impact of reduction of MRPL-3 and MRPS-22 by RNAi in wildtype and suppressor worms, as mutations in human homologs of MRPL-3 and MRPS-22 cause mitochondrial dysfunction and infant mortality^54,55^. Inhibition of *mrpl-3* or *mrps-22* by RNAi in wildtype worms delayed development. However, the rate of development of worms raised on *mrps-22*(RNAi) or *mrpl-3*(RNAi) was increased in *mrps-38(P210L)* worms and *mrps-15(A43V)* worms treated with *mrpl-3*(RNAi) Figure 6G).

Analysis of data from genome-wide CRISPR KO screens in a large variety of human cancer cell lines can give insight into gene function by scoring the response to perturbation of a gene and comparing amongst the dataset, which can be ranked by similarity using a Co-Dependency score^56^. Interestingly, the Top 100 DepMap Co-Dependency scores showed that CRISPR KO of human LONP1 is strongly correlated to CRISPR KO of genes that regulate protein synthesis on mitochondrial ribosomes (Figure 6H, Supplementary Table 6). The correlation between LONP1 and mitochondrial translation components in hundreds of human cell lines supports a conserved role for LONP-1 in the regulation of mitochondrial translation.

## Discussion

When LONP-1 is compromised, ATFS-1 accumulates in the mitochondrial matrix, promoting an increase in mtDNA replication^22^. The increase in mtDNA replication has a corresponding increase in mtRNA transcription, raising the levels of mt-rRNA, which initiates biogenesis of mitochondrial ribosomes, alongside increased levels of mt-tRNAs and mt-mRNA, which are all processed from polycistronic transcripts^57^. Simultaneously, degradation of imported, unassembled mitochondrial ribosomal proteins is reduced, which promotes their assembly into mitochondrial ribosomes. Increased mitochondrial ribosome abundance promotes mtDNA-encoded OXPHOS protein synthesis that is not in balance with the imported nuclear-encoded OXPHOS proteins, which can inhibit formation of functional OXPHOS complexes. Further, reduced chaperone activity of LONP-1 impairs OXPHOS assembly. The lack of degradation of unassembled OXPHOS proteins by LONP-1 would also obstruct OXPHOS biogenesis. This loss of OXPHOS causes mitochondrial stress, leading to developmental delay.

The finding that a mutation in the mitochondrial ribosome protein MRPS-38 restores OXPHOS in *lonp-1* mutant worms suggests that altering protein synthesis within the mitochondrial matrix can impact OXPHOS assembly. Our data suggest that the restoration of OXPHOS in *lonp-1* mutant worms by suppressor mutations likely occurs by reducing mitochondrial protein synthesis, as the highly abundant mitoribosomes do not appear to be translating at a wildtype rate. In this scenario, the slowing of mitochondrial protein translation enhances OXPHOS assembly. Direct inhibitors of mitochondrial translation do not suppress the mitochondrial dysfunction caused by inhibition of LONP-1 expression, suggesting that the reduction in translation by MRPS-38 is related to a context-specific function of the small subunit of the mitochondrial ribosome. The yeast homolog of MRPS-38, ms38, interacts with mt-mRNA and the peptidyl transfer center which can impact mitochondrial protein synthesis via multiple mechanisms^58^. Further studies on the *C. elegans* mitochondrial ribosome structure and mitoribosome footprinting may elucidate the exact mechanism by which this occurs.

Ultimately, our data suggest that the slowing of mitochondrial translation allows competent assembly of OXPHOS complexes, which is deficient in *lonp-1* mutant worms. Intriguingly, transcripts for a subset of mitochondrially-encoded complex I proteins are reduced in the *lonp-1(cmh24);mrps-38(P210L)* worms compared to *lonp-1(cmh24)* worms. Assembly of complex I has not been fully elucidated, however it has been suggested that formation of a complex containing ND2, ND3, ND4L and ND6, denoted as the PP-b module, is a required step in the biogenesis of complex I.^49^ The corresponding complex I proteins do not appear significantly reduced in *lonp-1(cmh24);mrps-38(P210L)* worms, suggesting that the reduction of mt-mRNAs does not reduce protein synthesis. Therefore, it is likely that the decrease in mRNA transcripts related to the PP-b module occurs post-translationally. Several assembly factors interact with the mitochondrial ribosome and nascent proteins of the PP-b module to form the subcomplex^59,60^. Slowing synthesis of nascent proteins may enhance this activity, or MRPS-38 may directly influence these interactions, leading to increased assembly of OXPHOS complexes. Our findings suggest that mtDNA-encoded proteins initiate OXPHOS complex biogenesis, highlighting a mechanism by which mitochondria can regulate OXPHOS independent from the nucleus. Further studies on *C. elegans* OXPHOS assembly at the mitochondrial ribosome may shed light on this process.

The increase in mitochondrial protein synthesis observed in the *lonp-1(cmh24)* mitochondria resembles the increase in translation observed in mitochondria harvested from the hearts of MTIF3 KO mice^61^. Further, MTIF3 KO mice exhibit reduced OXPHOS proteins and OXPHOS complex abundance similar to our observations in *lonp-1(cmh24)* mitochondria. We speculate that LONP-1 dysfunction causes a loss of regulation of mitochondrial translation initiation by MTIF3. MTIF3 may be regulated directly by LONP-1, as heart specific KO of LONP-1 in mice causes MTIF3 to become insoluble, or it may be indirectly affected by the disruption of the mitochondrial RNA granule, a phase separated hub of RNA processing proteins, several of which are LONP-1 substrates^34^. MTIF3 promotes dissociation of the mitochondrial ribosome^62^. Curiously, we did not observe tagged mitochondrial ribosome proteins MRPS17 or MRPL49 in sucrose gradient fractions from *lonp-1(cmh24)* mitochondria that correspond to unassembled mitoribosomes, suggesting that *lonp-1(cmh24)* monosomes are more stable. Dissociation of monosomes may cause altered mt-mRNA occupancy and degradation which may explain the reduction in mt-mRNAs in *lonp-1(cmh24);mrps-38(P210L)* compared to *lonp-1(cmh24)* worms. Structural analyses of MRPS38 and MRPS15 homologs suggest interactions that could influence mitoribosome-bound MTIF3. Further studies are necessary to determine how MTIF3 promotes dissociation of the mitochondrial ribosome and how translation initiation from monosomes may alter mRNA dynamics.

In bacteria, the Lon protease has been shown to degrade several different ribosomal proteins^63^. Replication factors have also been identified as substrates of bacterial Lon^63^. The role of Lon as a “master regulator” appears conserved from bacteria to humans^64^. Lastly, analysis of DepMap data indicates that inactivation of LONP-1 correlates with loss of factors involved in mitochondrial translation in hundreds of human cell lines^56^. We therefore propose that the role of LONP-1 in degrading mitochondrial ribosomal proteins to regulate mitochondrial translation is conserved from bacteria to eukaryotic mitochondria.

This study highlights the critical role of mitochondrial translation in maintaining respiratory function. We demonstrate that disruption of proteostasis caused by LONP-1 impairment perturbs OXPHOS, which can be rescued by altering mitoribosome function. Notably, Huntington’s disease cells were recently reported to display altered mitochondrial ribosome occupancy, suggesting that mitoribosome dysregulation may be a common feature of both inherited mitochondrial disorders and neurodegenerative disease^65^. These findings raise the possibility that targeted modulation of mitochondrial translation could offer a therapeutic strategy across diverse pathological contexts.

## Materials and Methods

### Worm strains

Worms were raised on NGM plates and fed HT115 *Escherichia coli*. The N2 strain provided by the *Caenorhabditis* Genetics Center (Minneapolis, Minnesota) was used as wildtype in all experiments. The strains *lonp-1(cmh24), lonp-1(cmh25), mrps-15(A43V), mrps-38(S42L), mrps-38(P210L), lonp-1(R711G), spg-7(M666G), mrpl-58(flag), mrps-17(ollas)* were generated using CRISPR-cas9 as previously described, with reagents donated by the Mello laboratory^66^. Double mutants of *lonp-1(cmh24); mrps-15(A43V)* and *mrpl-58(flag); mrps-17(ollas)* were generated using two CRISPR-cas9 edits. All CRISPR-cas9 edited strains were outcrossed 3 times. Guide and donor sequences are provided on Supplementary Table.

### Western Blot

Worms were bleach synchronized, raised on NGM plates at 20°C, fed HT115 *Escherichia coli*, harvested at the L4 stage and washed three times in S-basal. Worm pellets were flash frozen and lysed in detergent buffer using a Teflon homogenizer or BeadBlaster homogenizer. Commercial antibodies against TUBULIN (1:1000 BioRad MCA78G), OLLAS (1:1000 Thermo MA5-16125), FLAG (1:1000 Sigma F1804), MTCO1 (1:1000 Abcam AB14705). Previously validated custom antibodies against LONP-1, POLG-1 and ATFS-1 were used^22,67^. Blots were imaged using ChemiDoc XRS.

### Development Assay

Worms were bleach synchronized, fed HT115 *Escherichia coli* and raised at 20°C. Plates were analyzed by microscopy and animals were scored by stage after 72 hours. At least 50 animals were scored per replicate. Experiment was performed in triplicate. The number of animals at the L4 and adult stages was summed and divided by the total number of animals to give the percentage.

### Mitochondrial Isolation

Mitochondrial isolation was performed as described previously^68^. Worms were bleach synchronized, fed HT115 *Escherichia coli,* raised on NGM plates at 20°C, harvested at the L4 or D12 adult stage and washed three times in S-basal. Worm pellets were flash frozen and lysed in detergent buffer using a Teflon homogenizer or BeadBlaster homogenizer. L4 or D1 adult worm pellets greater than 100 μL were resuspended in Mitochondrial buffer (70 mM sucrose, 1 mM EGTA, 210 mM sorbitol, 10 mM MOPS (pH 7.4)) equivalent to two-fold the pellet volume then lysed using either a Teflon homogenizer or BeadBlaster. Lysates were submitted to differential centrifugation (450g 10min;550g 10min; 9500g 10min) then resuspended in 1 mL of buffer then centrifuged again (500g 10min, 9500g 10 min) to create a pellet enriched for mitochondria. 5 μL were then treated with detergent and measured by Bradford assay for protein estimation.

### Blue Native Page In-Gel Assay

Assay was performed as previously described^69^. Worms were bleach synchronized, fed HT115 *Escherichia coli*, raised on NGM plates at 20°C, harvested at the Day 1 adult stage and washed three times in S-basal. Worm pellets were flash frozen and lysed in detergent buffer using a Teflon homogenizer or BeadBlaster homogenizer. Mitochondria were isolated from synchronized Day 1 adult animals. 200 μg of mitochondrial protein (determined by Bradford assay) was pelleted and incubated for 10 minutes at room temperature in a mixture containing 5 μL of 4X sample buffer, 6 μL of 6igitonin, and 9 uL of water. The mixture was then centrifuged for 20 minutes at 15,000 RPM. The supernatant was transferred to a new tube and 2 μL of 5% NativePAGE G-250 Coomassie was added. The sample mix was then loaded into an Invitrogen NativePAGE 3-12% Bis-Tris gel and run at 100V until all samples entered the gel, at which point voltage was increased to 300V. After samples ran through one third of the gel, the buffer was replaced with cathode buffer (10-fold diluted G-250 Invitrogen Running Buffer kit). After running, the gel was incubated in 10mL complex I buffer (41mg Nitroblue tetrazolium, 78.8mg TrisHCl, 35.5 mg NADH in 10mL) for 40 minutes to 3 hours. One lane containing NativeMark protein standard was excised and incubated in Colloidal Coomassie stain to visualize the ladder bands.

### Thrash Assay

Assay was performed as previously described^70^. Worms were bleach synchronized, fed HT115 *Escherichia coli* and raised on NGM plates at 20°C until reaching the the Day 1 adult stage. Adult worms were transferred into a droplet of S Basal and after waiting 1 minute body bends were counted and recorded for 30 seconds. 10 worms were assayed for each replicate.

### Microscopy – TMRE

Imaging was performed as previously described^71^. Worms were bleach synchronized, fed HT115 *Escherichia coli* and raised on NGM plates at 20°C until reaching the L4 stage. L4 worms were placed into 1mL of 10 μM TMRE in S Basal for 45 minutes at room temperature while agitated on a shaker. After TMRE incubation worms were transferred to NGM plates and allowed to recover for 3 hours. L4 worms were then transferred to a slide with an agarose pad containing 3 μL of levamisole. The Zeiss LSM800 microscope with Airyscan was used to image fluorescence of the intestinal mitochondria at 63X magnification.

### Quantification of mtDNA

mtDNA quantification was performed as previously described^72^. Worms were bleach synchronized, fed HT115 *Escherichia coli* and raised on NGM plates at 20°C until reaching the L4 stage. 25 worms were collected in 25 μL of Lysis Buffer (50 mM KCl, 10 mM Tris-HCl (pH 8.3), 2.5 mM MgCl_2_, 0.45% NP-40, 0.45% Tween 20, 0.01% gelatin, with freshly added 200 μg ml^−1^ proteinase K) and frozen, then submitted to a thermocycler lysis protocol (60 C 65 min; 95 C 15 min). The lysate was then diluted to 100 μL with water. 2uL of lysate was used per reaction in triplicate, with a set of primers specific to mtDNA and a set of primers for ges-1, an intestinal cell transcription factor used for normalization. Each reaction contained 5 uL Thermo Scientific SyBr Green Maxima Mix, 1 uL primer mix and 2 uL water. qPCR was performed on the MyiQ2 Two-Color Real-Time PCR Detection System (Bio-Rad Laboratories).

### S^35^ Translation Assay

Analysis was performed as previously described^73^. Worms were bleach synchronized, fed HT115 *Escherichia coli* and raised on NGM plates at 20°C until reaching the Day 1 adult stage. Mitochondria were isolated from D1 adults. 200 ug of isolated mitochondria was incubated in translation mix (25 mM sucrose, 75 mM sorbitol, 100 mM KCl, 1 mM MgCl_2_, 0.05 mM EDTA, 10 mM Tris–HCl and 10 mM K_2_HPO_4_, pH 7.4) containing 10 mM glutamate, 2.5 mM malate, 1 mM ADP, 1 mg/ml fatty acid-free BSA, 100 μg/ml emetine, 100 μg/ml cycloheximide and 0.2 mM amino acids (minus cysteine and methionine (Promega). The translation mixture (95 μl) was pre-warmed at 20 °C for 5 min. Next 5 μl of EasyTag™ EXPRESS^35^S Protein Labeling Mix, [^35^S]-, 2mCi was added and translation was allowed to proceed for 60 min at 20 °C. The mix was split and one half was treated in parallel with 200 μM chloramphenicol after 5 minutes of translation as a control. Mitochondria were then washed three times with MSEM buffer containing 200 mM cysteine and methionine. SDS-Page was performed and gels were fixed and stained with Coomassie, then imaged. Gels were then vacuum dried and exposed to a Phosphor screen for 72 hours before being visualized using a Typhoon imager.

### Mass Spectrometry

Mass spectrometry was performed by the Thermo Fisher Center for Multiplexed Proteomics in the Department of Cell Biology at Harvard Medical School. Worms were bleach synchronized, fed HT115 *Escherichia coli* and raised on NGM plates at 20°C until reaching the Day 1 adult stage. Mitochondria were isolated and 100 μg of protein was submitted for proteomics. 25 μg of protein from each sample was reduced with TCEP, alkylated with iodoacetamide, and then further reduced with DTT. Proteins were precipitated onto SP3 beads to facilitate a buffer exchange into digestion buffer. Samples were digested with Lys-C (1:50) overnight at room temperature and trypsin (1:50) for 6 hours at 37°C. Peptides were labelled with TMT 18plex reagents. 2 μL of each sample was pooled and used to shoot a ratio check in order to confirm complete TMT labelling and to allow for normalization of each sample. All 16 TMTPro-labelled samples were pooled according to the ratios determined from the ratio check. Peptides were desalted using a Sep-pak. Peptides were fractionated into 24 fractions using basic reverse phase HPLC and 6 fractions were solubilized, desalted by stage tip, and analyzed on an Orbitrap Eclipse mass spectrometer with a FAIMS device enabled. *C. elegans* mitochondrial proteins were inferred from previous experiments and MitoCarta homology^27,36,37^. Peptide spectral matches were filtered to a 1% false discovery rate (FDR) using the target-decoy strategy combined with linear discriminant analysis. Proteins were quantified only from peptides with a summed SN threshold of >100. Abundance of mitochondrial proteins were compared to wildtype to determine fold changes. Genes with log2foldchange>0.15 were classfied enriched, log2foldchange<-0.15 were classified depleted. Lists of all enriched and depleted proteins with were submitted to STRING db for GO analysis^74^. The processed proteomics dataset is available at Zenodo with the following DOI: https://doi.org/10.5281/zenodo.18187177

### RNA-sequencing

Worms were bleach synchronized, fed HT115 *Escherichia coli* and raised on NGM plates at 20°C until reaching the L4 stage. Pellets were washed 3 times in S-basal then flash frozen in liquid nitrogen. RNA was isolated using Trizol and Phenol-Chloroform extraction. Total RNA was sent to Innomics which performed library preparation and 100 bp paired-end mRNA-sequencing by DNBseq. QC and analysis was done using OneStopRNAseq (https://mccb.umassmed.edu/OneStopRNAseq/). RNA reads were aligned to the WS295 reference genome, and differential gene expression analysis was performed with edgeR43. Reads of all transcripts were compared to wildtype to determine fold changes. Genes with log2foldchange>0.5 were classfied upregulated, log2foldchange<-0.5 were classified downregulated. Lists of all up and downregulated genes with FDR>0.5 were submitted to STRING db for GO analysis^74^. Sequencing data are available at Gene Expression Omnibus (GEO) session GSE315983.

### Mutagenesis screen

Approximately 20,000 worms were bleached and raised on plates until reaching the L4 stage. Worms were treated with 50mM EMS (ethyl methanesulfonate). Gravid animals (approx. 10 eggs/worm) were bleached to give approximately 200,000 mutagenized haploid genomes in the F1 generation. F1 worms were bleached to 40 large plates for F2 homozygous mutants and plates were screened for worms with improved development. Selected strains were outcrossed 3X to select for phenotype. Genomic DNA was isolated from candidate suppressor worms using Qiagen Gentra Puregene Tissue Kit. 100bp PE Whole genome sequencing with 5GB sequencing depth and library preparation was performed by BGI.

### Respiration Assay

The Seahorse Extracellular Flux Analyzer XFe96 (Seahorse Biosciences) assay was performed as previously described^72^. Worms were bleach synchronized and cultured to the L4 stage. Worms were transferred into 10 wells of a 96-well microplate containing 180 μl M9 buffer with 10 worms/well. Basal respiration was measured for a total of 30 min, in 6 min intervals that included a 2 min mix, a 2 min time delay, and a 2 min measurement. To measure respiratory capacity, 15 μM carbonyl cyanide-4-(trifluoromethoxy)phenylhydrazone was injected, the oxygen consumption rate reading was allowed to stabilize for 6 min and then measured for five consecutive intervals. Mitochondrial respiration was blocked by adding 40 mM sodium azide.

### Sucrose Gradient Fractionation

Sucrose fractionation was performed as previously described^75^. Worms were bleach synchronized, fed HT115 *Escherichia coli* and raised on NGM plates at 20°C until reaching the Day 1 adult stage. Mitochondria were isolated as described above and 200 μg of mitochondrial protein was pelleted, then resuspended in 200 μL Lysis buffer containing 3% (w/v) sucrose, 100 mM NH_4_Cl, 15 mM MgCl_2_, 20 mM Tris-HCl pH 7.5, cOmplete protease inhibitor cocktail (Roche) and 0.08 U per µl RiboLock RNase Inhibitor (Thermo Fisher)) for 30 min at 4 °C with gentle shaking and the resulting lysate was cleared at 16,000*g* for 15 min at 4 °C before loading onto a sucrose gradient (7.5–30% (w/v) sucrose, 100 mM NH_4_Cl, 10 mM MgCl_2_, 20 mM Tris-HCl pH 7.5 and cOmplete protease inhibitor cocktail (Roche)). Lysates were spun by ultracentrifugation for 15 h at 158,000*g* (30,400 r.p.m.; high-resolution gradient) using an SW41Ti rotor (Beckman Coulter). 18 fractions were collected by pipetting 450 μL into tubes, and protein was precipitated using TCA. Protein pellets were resuspended in sample buffer and submitted to SDS-page and western blot.

### Mitochondrial Ribosome Inhibition Assay

1.5 mg/mL doxycycline hydroxychloride (Sigma D447) stock solution was added to 300 μL S-basal and then pipetted onto NGM plates seeded with HT115 *Escherichia coli*, then dried for 10 minutes. Worms were bleach synchronized and imaged every 24 hours to monitor growth.

### Protein Structural Analysis

*C. elegans mrps-38* inferred protein sequence was compared to protein sequences from *T. brucei* and *T. cruzi* using ClustalO^76^. Mitoribosome images were generated from PDB IDs 6ZM5, 7HIW and 7AOR structures using Mol*^77,78^.

### Gene Ontology Analysis of CRISPR DepMap

The Top 100 CRISPR Co-dependencies for LONP1 were generated by DepMap^56^. STRING GO analysis of all 100 genes was used to determine enriched biological processes^74^.

## Data availability

RNA-seq data supporting this study are openly available from the Gene Expression Omnibus (GEO) under accession number **GSE315983**. Proteomics data supporting this study are openly available at Zenodo DOI https://doi.org/10.5281/zenodo.18187177.

## Acknowledgments

We thank the *Caenorhabditis* Genetics Center for providing *C. elegans* strains (funded by NIH Office of Research 362 Infrastructure Programs (P40OD010440), and the UMass Medical School Core facilities for deep sequencing. This work was supported by the National Institutes of Health grants (R01AG040061 and R01AG047182) to C.M.H. The authors are solely responsible for the content. We would like to acknowledge Gyan Prakash for helpful discussions and suggestions, and Thermo Fisher Scientific Center for Multiplexed Proteomics at Harvard Medical School (https://tcmp.hms.harvard.edu). We are grateful to Nicholas Petersen and Samantha Tse for help with the mutagenesis screen.

## Author contributions

L.A. and C.M.H. planned the experiments. L.A. and Y.D. generated worm strains and western blots. A.M. performed RNA-seq. R.L., and L.J.Z. analyzed the sequencing data. L.A. performed all other experiments. L.A. and C.M.H. wrote the manuscript.

---

**Figure S1.**
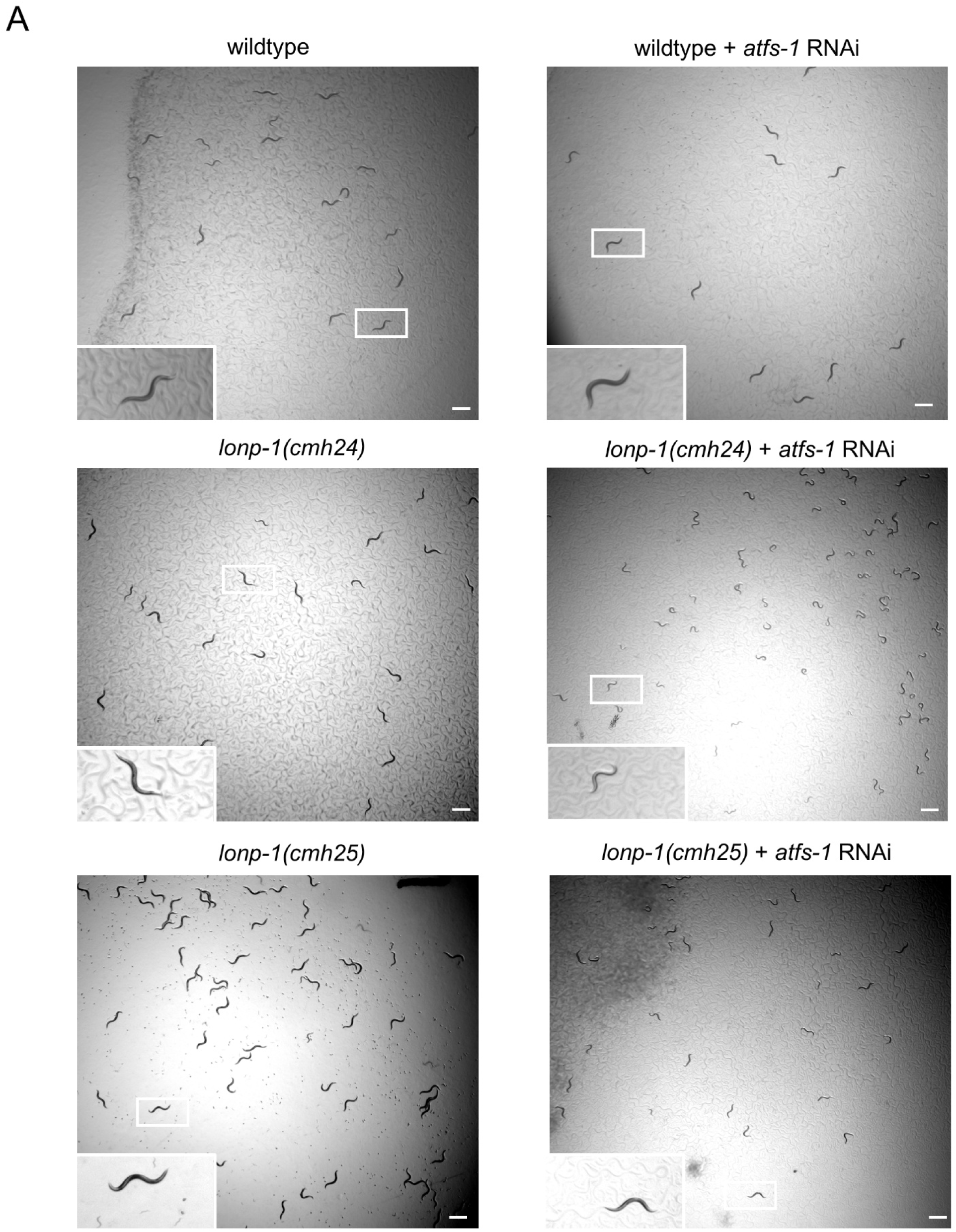
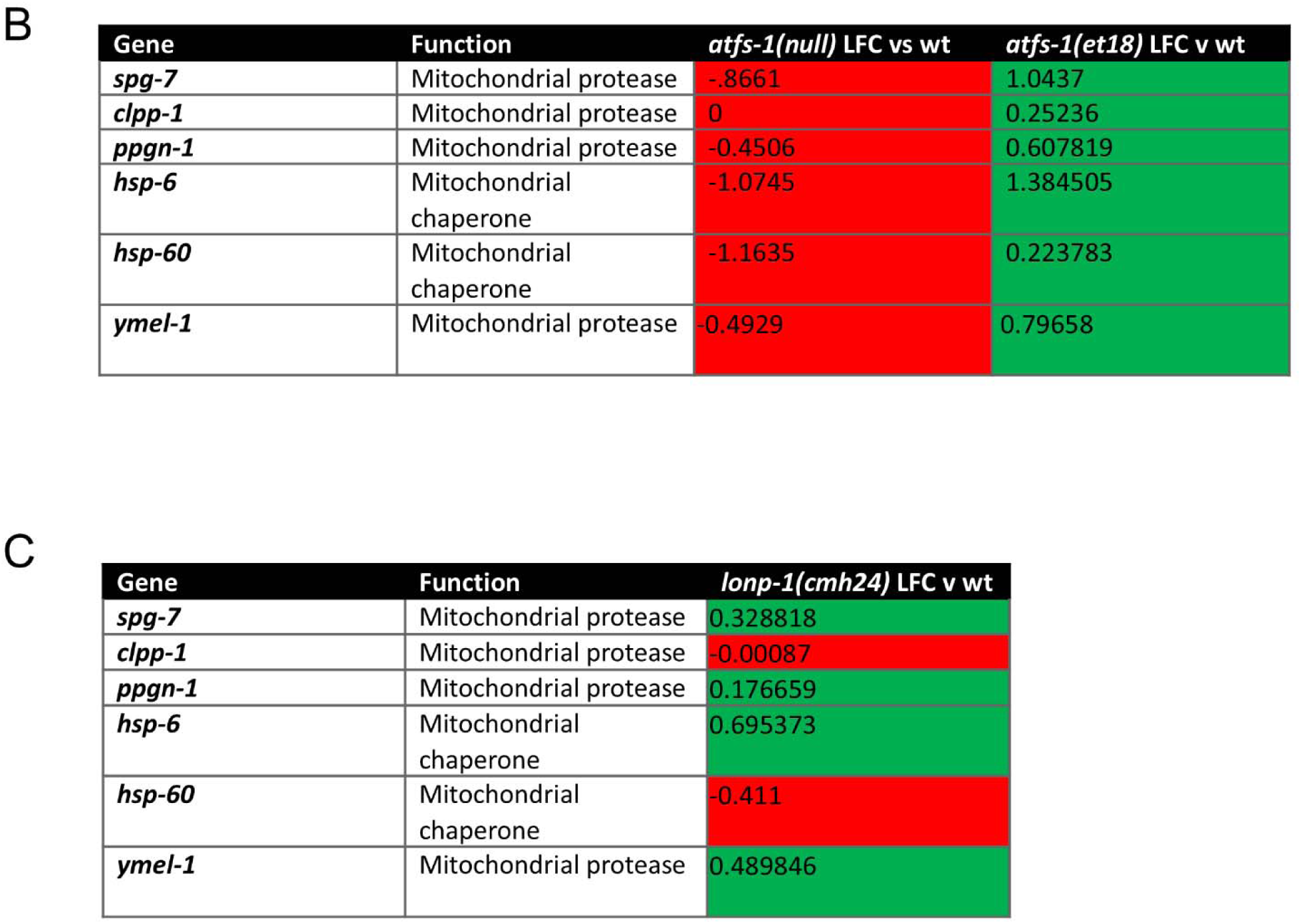
*atfs-1* induces a protective response during *lonp-1* dysfunction. **(A)** Representative brightfield images of wildtype, *lonp-1(cmh24)* and *lonp-1(cmh25)* worms after 72 hours growth on *atfs-1* RNAi at 11.3X magnification (scale bar, 1 mm). **(B)** log2foldchange values from RNA-seq from *atfs-1(null)* and *atfs-1(et18)* worms compared to wildtype worms for genes related to mitochondrial proteostasis, red values indicate decreased expression, green values indicate increased expression **(C)** log2foldchange values from RNA-seq from *lonp-1(cmh24)* worms compared to wildtype worms for genes related to mitochondrial proteostasis, red values indicate decreased expression, green values indicate increased expression. All depicted experiments had three biological replicates with similar results.

**Figure S2.**
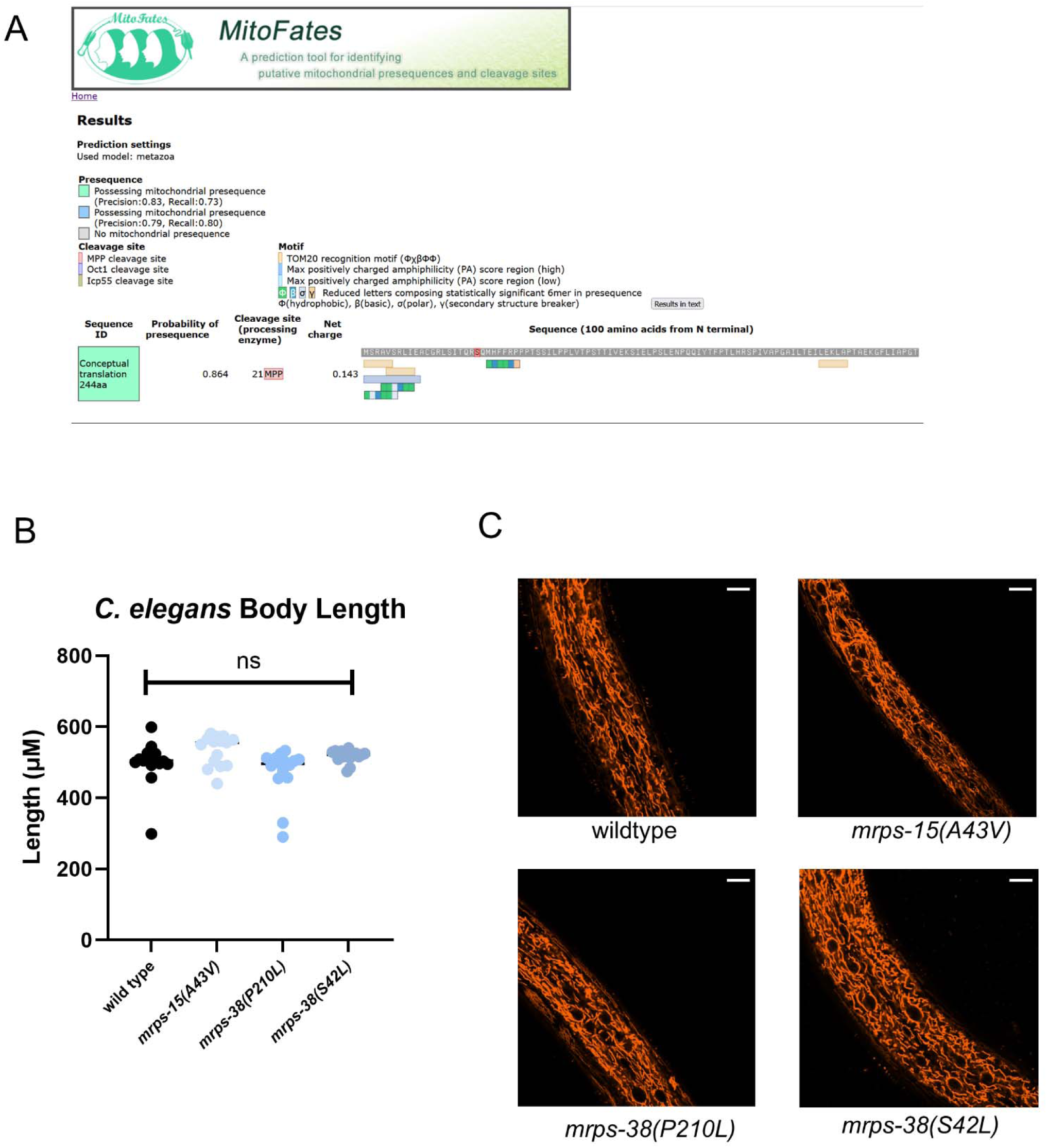
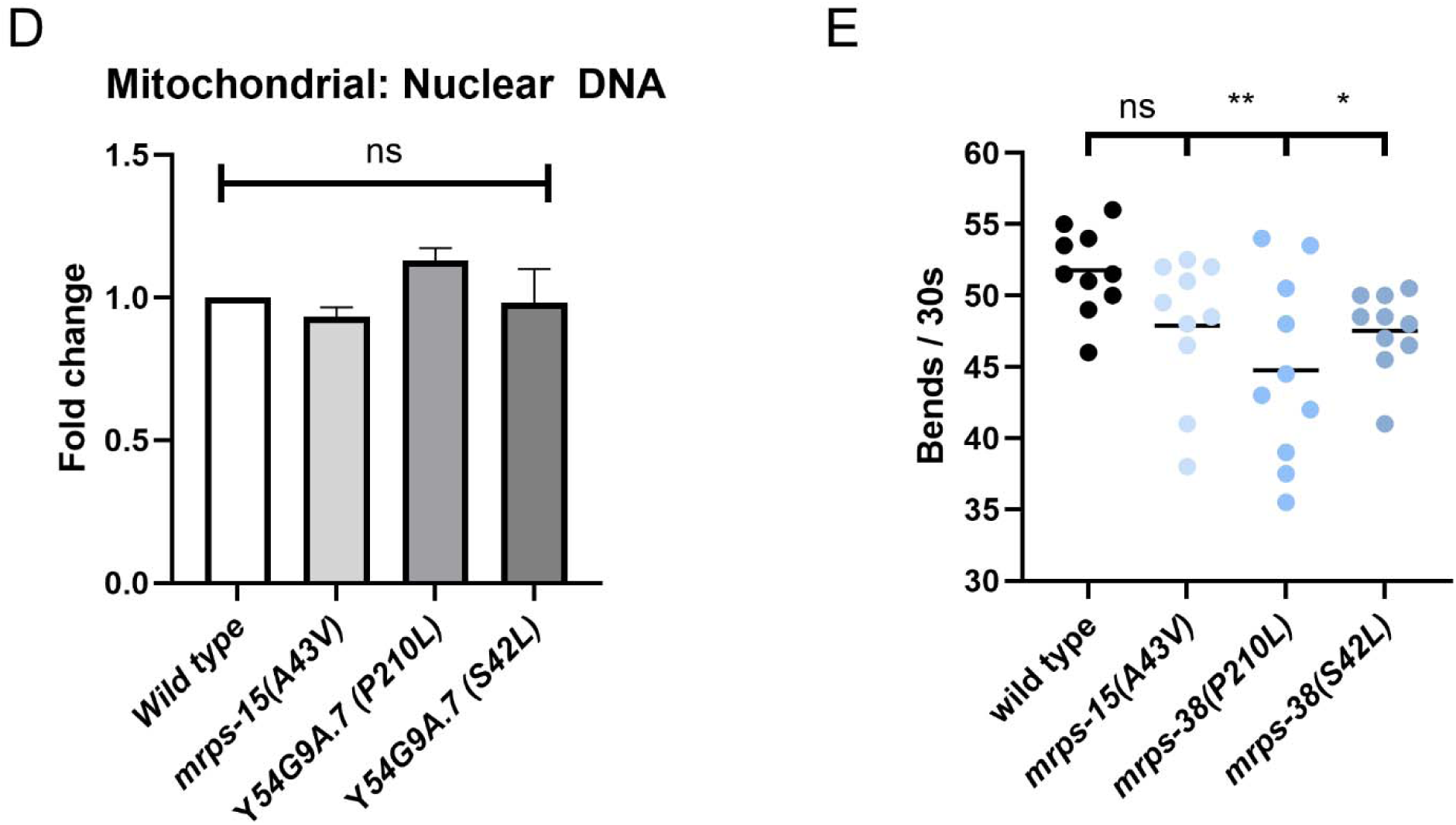
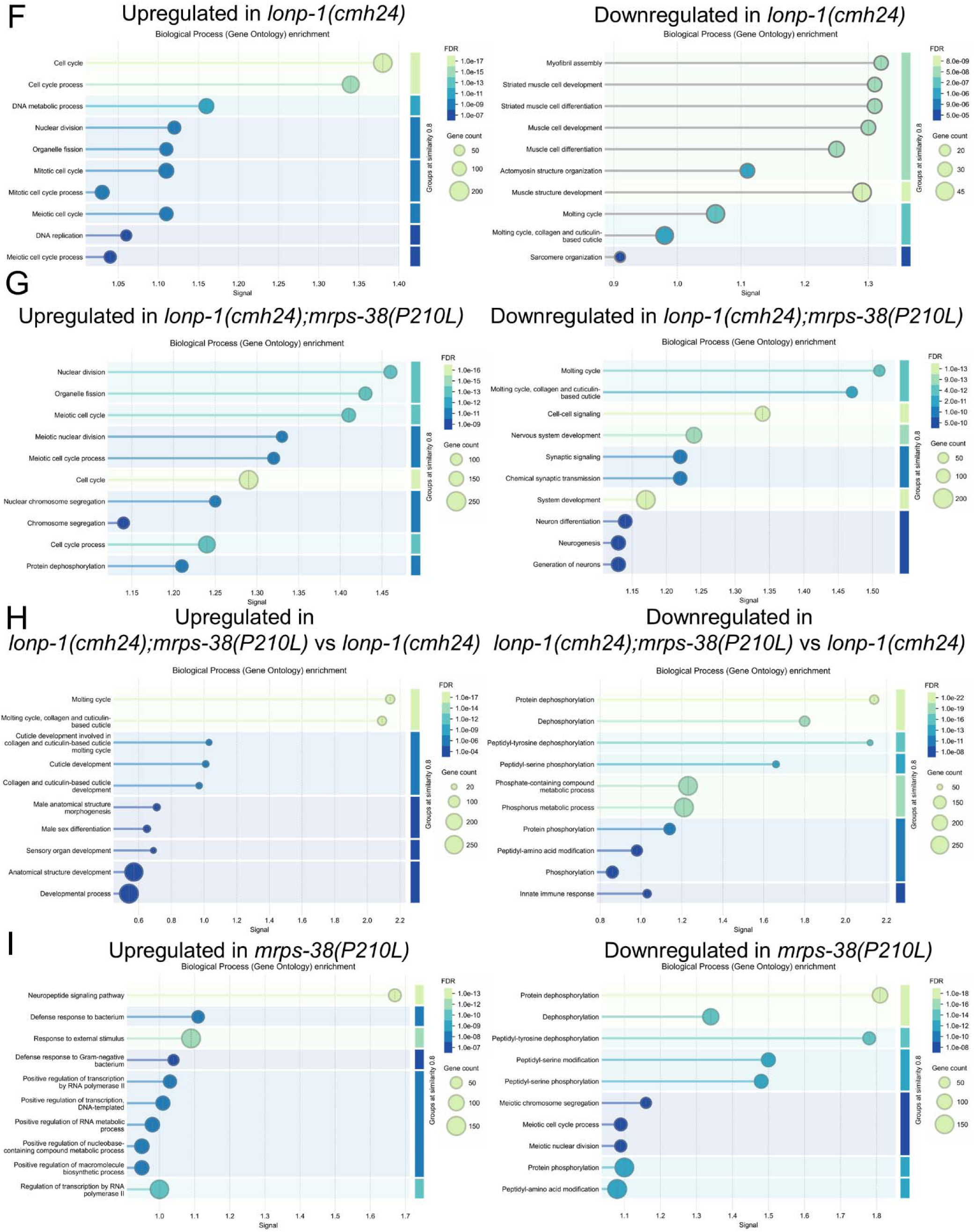
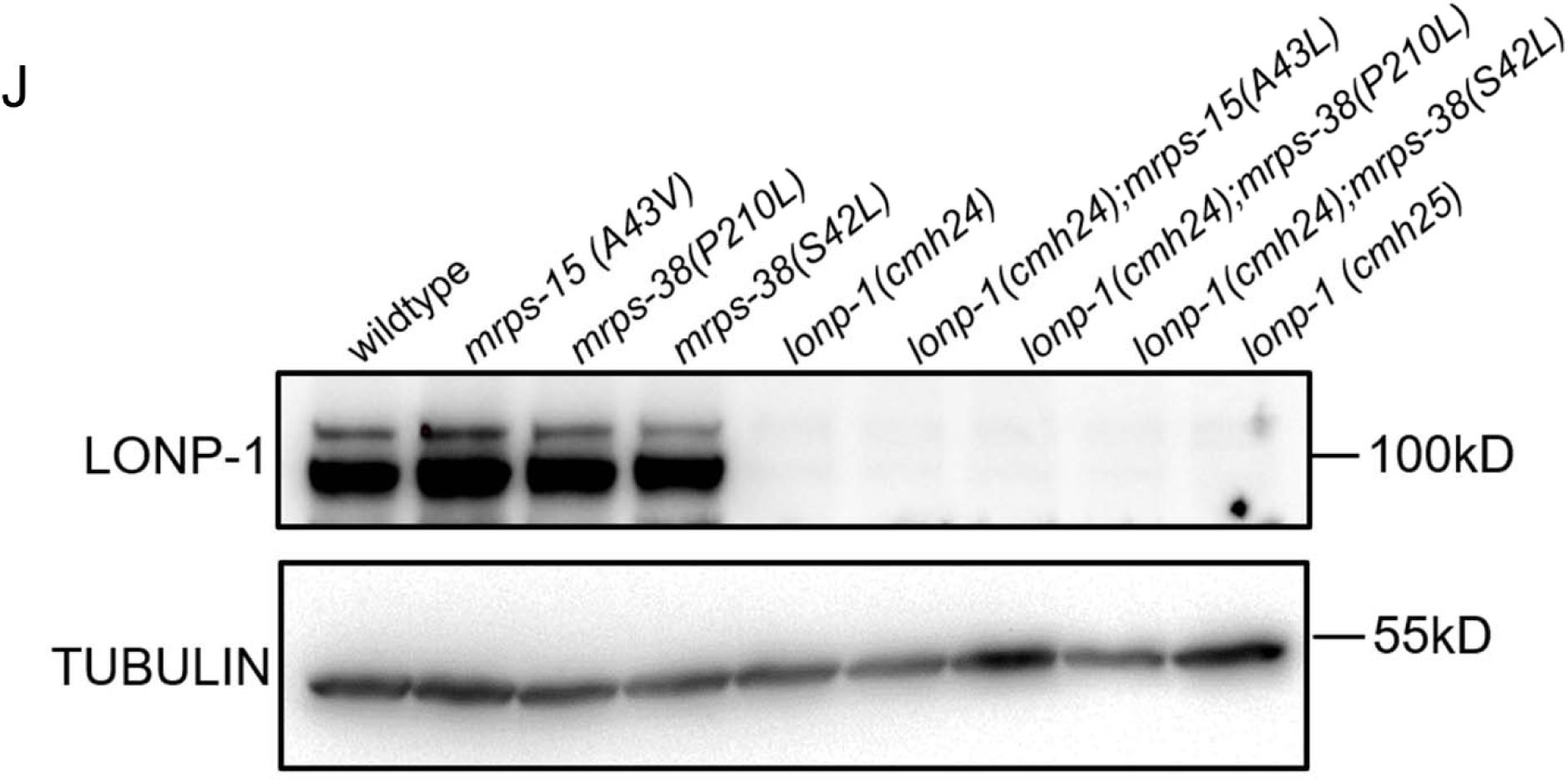
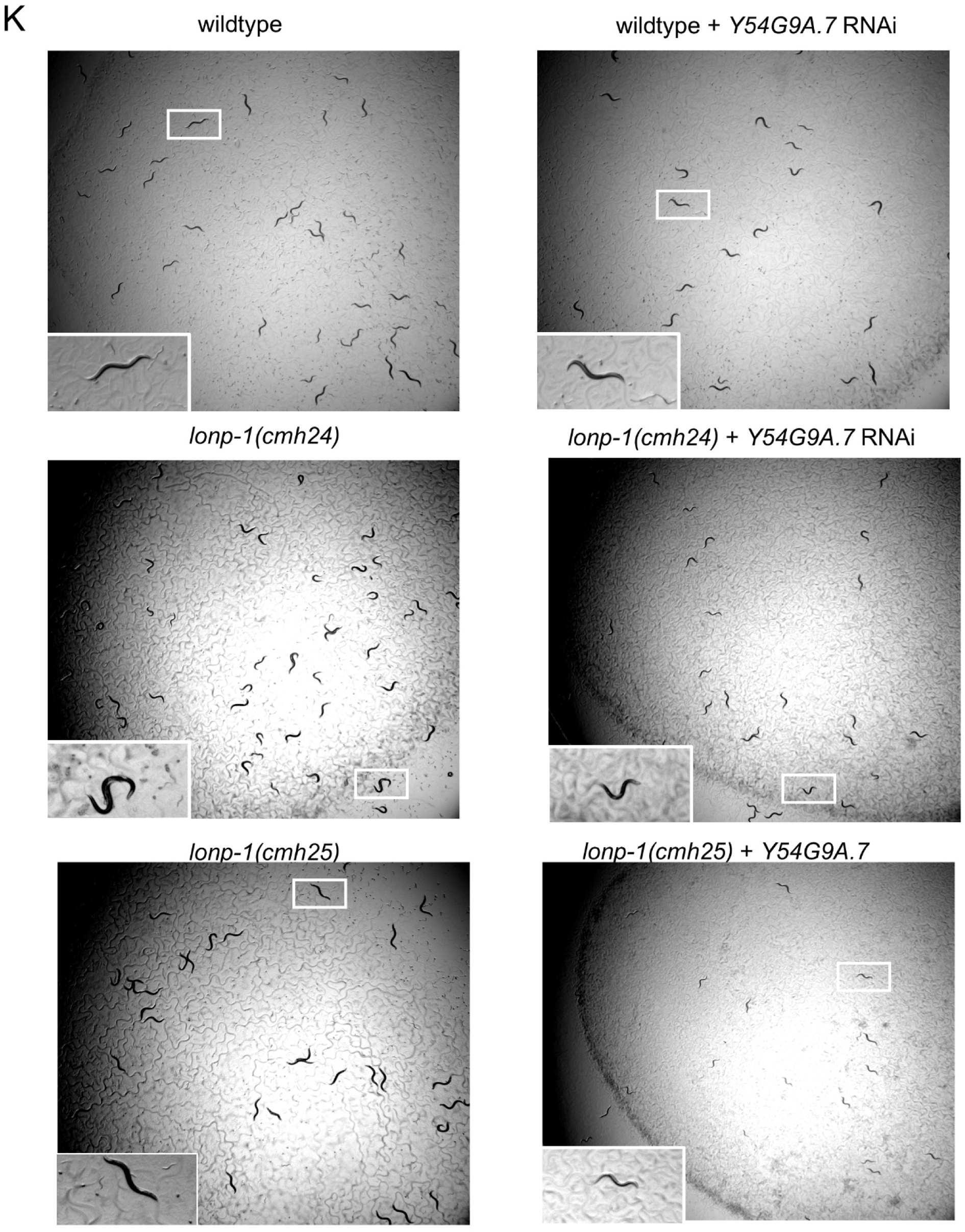
Characterization of mrps-38(P210L), mrps-38(S42L) and mrps-15(A43V) mitochondrial ribosome mutations. **(A)** MitoFates^81^ prediction of mitochondrial targeting sequence for Y54G9A.7 **(B)** Quantification of body length of wildtype, *mrps-15(A43V), mrps-38(P210L) and mrps-38(S42L)* worms after 72 hours of growth, each point is a single animal, n=30 NS, not significant (one-way ANOVA followed by Dunnett’s multiple comparison test) **(C)** Representative images of fluorescence of wildtype, *mrps-15(A43V), mrps-38(P210L) and mrps-38(S42L)* after TMRE staining at 63X magnification. Scale bar 10 µm **(D)** Quantification of mtDNA by qPCR of wildtype, *mrps-15(A43V), mrps-38(P210L) and mrps-38(S42L)* worms, n=3 biological replicates of 25 worms. Error bars mean ± SD (two-tailed Student’s *t*-test NS, not significant **(E)** Thrash assay of wildtype, *mrps-15(A43V), mrps-38(P210L) and mrps-38(S42L*) worms where each point is the number of body bends measured in an individual animal after 30 seconds, n=10 *p<0.05 **p<0.01 NS, not significant (two-way ANOVA with post-hoc Sidak’s test). **(F)** Left: STRING GO^74^ analysis of genes with adjusted shrunken log2foldchange>0.585 vs wildtype from RNA-seq of *lonp-1(cmh24)* worms. Right: STRING GO analysis of genes with adjusted shrunken log2foldchange<-0.585 vs wildtype from RNA-seq of *lonp-1(cmh24)* worms. FDR 0.05 **(G)** Left: STRING GO analysis of genes with adjusted shrunken log2foldchange>0.585 vs wildtype from RNA-seq of *lonp-1(cmh24);mrps-38(P210L)* worms. Right: STRING GO analysis of genes with adjusted shrunken log2foldchange<-0.585 vs wildtype from RNA-seq of *lonp-1(cmh24);mrps-38(P210L)* worms. FDR 0.05 **(H)** Left: STRING GO analysis of genes with adjusted shrunken log2foldchange>0.585 vs wildtype from RNA-seq of *mrps-38(P210L)* worms. Right: STRING GO analysis of genes with adjusted shrunken log2foldchange<-0.585 vs wildtype from RNA-seq of *mrps-38(P210L)* worms. **(I)** Left: STRING GO analysis of genes with adjusted shrunken log2foldchange>0.585 vs *lonp-1(cmh24)* from RNA-seq of *lonp-1(cmh24);mrps-38(P210L)* worms. Right: STRING GO analysis of genes with adjusted shrunken log2foldchange<-0.585 vs *lonp-1(cmh24)* from RNA-seq of *lonp-1(cmh24);mrps-38(P210L)* worms. **(J)** Western blot of wildtype, *lonp-1* mutants with suppressors for LONP-1 and TUBULIN **(K)** Representative brightfield images of wildtype, *lonp-1(cmh24)* and *lonp-1(cmh25)* worms after 72 hours growth (wildtype) or 120 hours growth (*lonp-1* mutants) on *Y54G9A.7* RNAi at 11.3X magnification (scale bar, 1 mm). All depicted experiments had three biological replicates with similar results.

**Figure S4.**
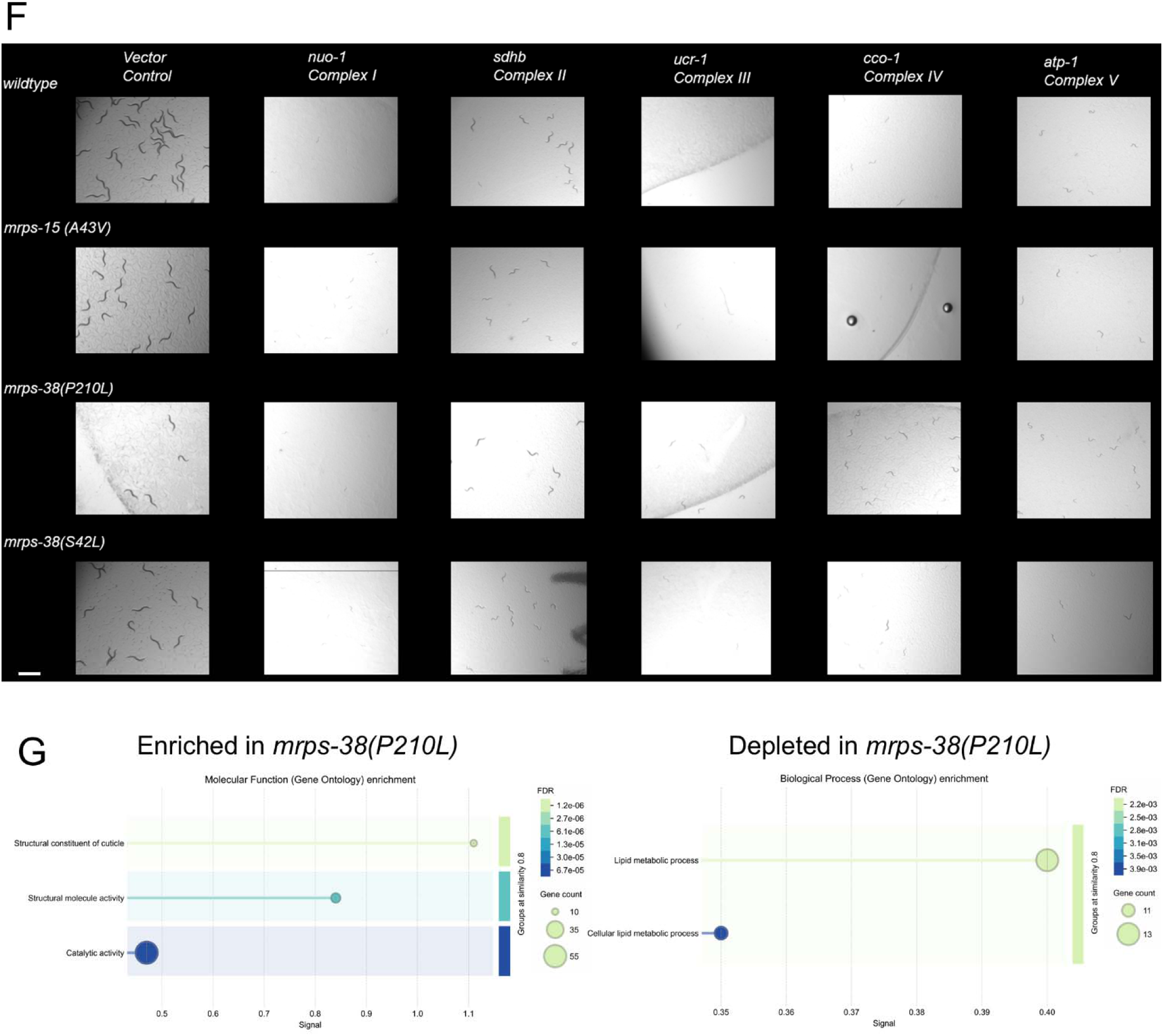
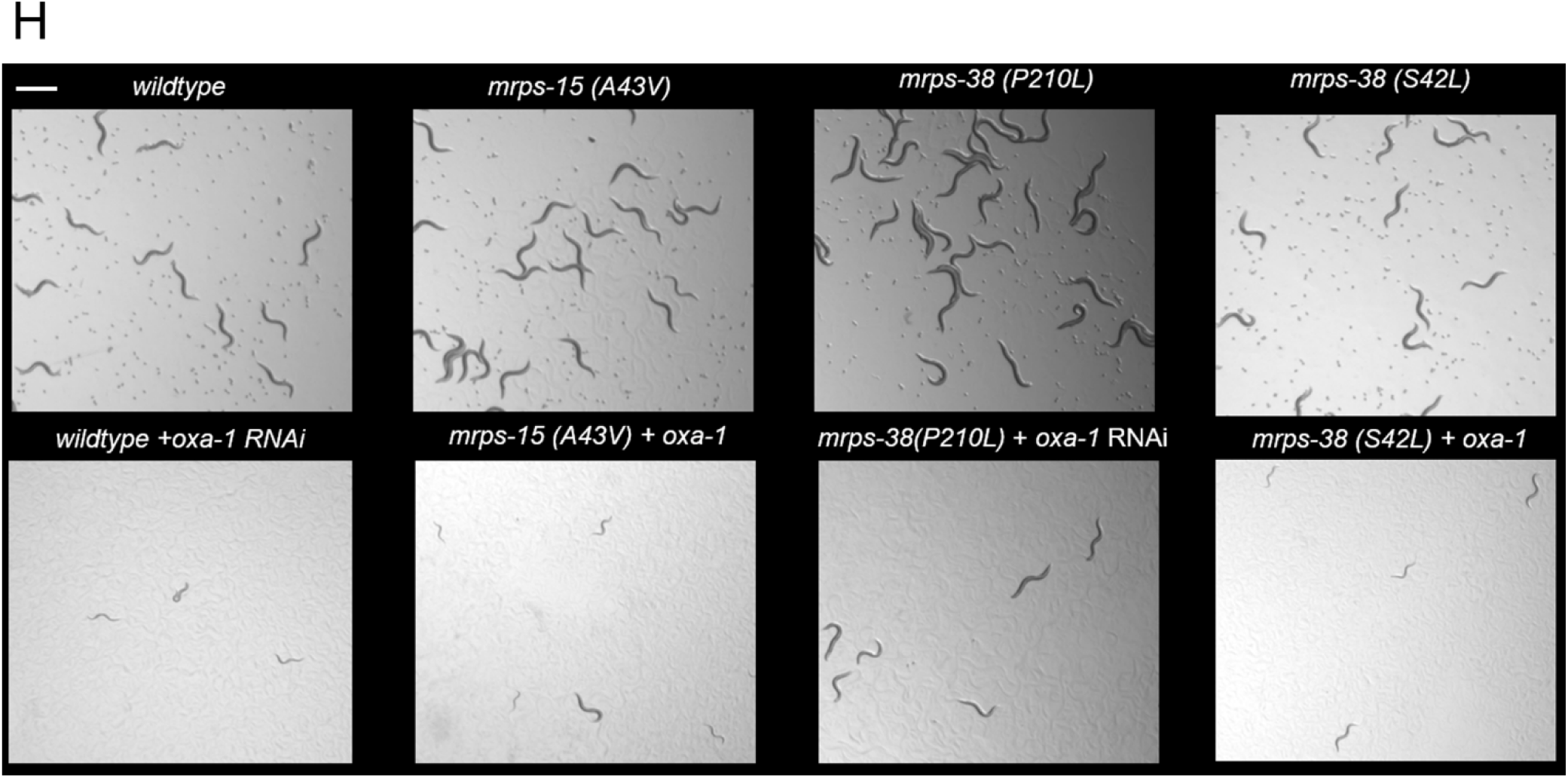
mrps-38(P210L), mrps-38(S42L) and mrps-15(A43V) restore impaired OXPHOS in lonp-1(cmh24) worms. **(A)** Left: SDS-Page followed by western blot of wildtype, *lonp-1(cmh24), mrps-15(A43V);lonp-1(cmh24)* and *mrps-15(A43V)* worm lysates for MT-COI and TUBULIN. Right: Quantification of western blot of MT-COI, n=3 biological replicates. Error bars mean ± SD (two-tailed Student’s *t*-test) *p<0.05 **(B)** Left: SDS-Page followed by western blot of wildtype, *lonp-1(cmh24), mrps-38(S42L);lonp-1(cmh24)* and *mrps-38(S42L)* worm lysates for MT-COI and TUBULIN. Right: Quantification of western blot of MT-COI, n=3 biological replicates. Error bars mean ± SD (two-tailed Student’s *t*-test) *p<0.05 **(C)** Left: STRING GO analysis of genes with log2foldchange>0.15 vs wildtype from mass spectrometry of mitochondria isolated from *lonp-1(cmh24)* worms. Right: STRING GO analysis of genes with log2foldchange<-0.15 vs wildtype from mass spectrometry of mitochondria isolated from *lonp-1(cmh24)* worms. **(D)** Left: STRING GO analysis of genes with log2foldchange>0.15 vs wildtype from mass spectrometry of mitochondria isolated from *lonp-1(cmh24);mrps-38(P210L)* worms. Right: STRING GO analysis of genes with log2foldchange<-0.15 vs wildtype from mass spectrometry of mitochondria isolated from *lonp-1(cmh24);mrps-38(P210L)* worms **(E)** Left: STRING GO analysis of genes with log2foldchange>0.15 vs wildtype from mass spectrometry of mitochondria isolated from *mrps-38(P210L)* worms. Right: STRING GO analysis of genes with log2foldchange<-0.15 vs wildtype from mass spectrometry of mitochondria isolated from *mrps-38(P210L)* worms **(F)** Representative brightfield images of wildtype, *mrps-15(A43V), mrps-38(P210L) and mrps-38(S42L*) worms after 72 hours growth on OXPHOS RNAi at 11.3X magnification (scale bar, 1 mm). **(G)** Left: STRING GO analysis of genes with log2foldchange>0.15 vs *lonp-1(cmh24)* from mass spectrometry of mitochondria isolated from *lonp-1(cmh24);mrps-38(P210L)* worms. Right: STRING GO analysis of genes with log2foldchange<-0.15 vs *lonp-1(cmh24)* from mass spectrometry of mitochondria isolated from *lonp-1(cmh24);mrps-38(P210L)* worms. **(H)** Representative brightfield images of wildtype, *mrps-15(A43V), mrps-38(P210L) and mrps-38(S42L*) worms after 72 hours growth on *oxa-1* RNAi at 11.3X magnification (scale bar, 1 mm). All depicted experiments had three biological replicates with similar results.

**Figure S5.**
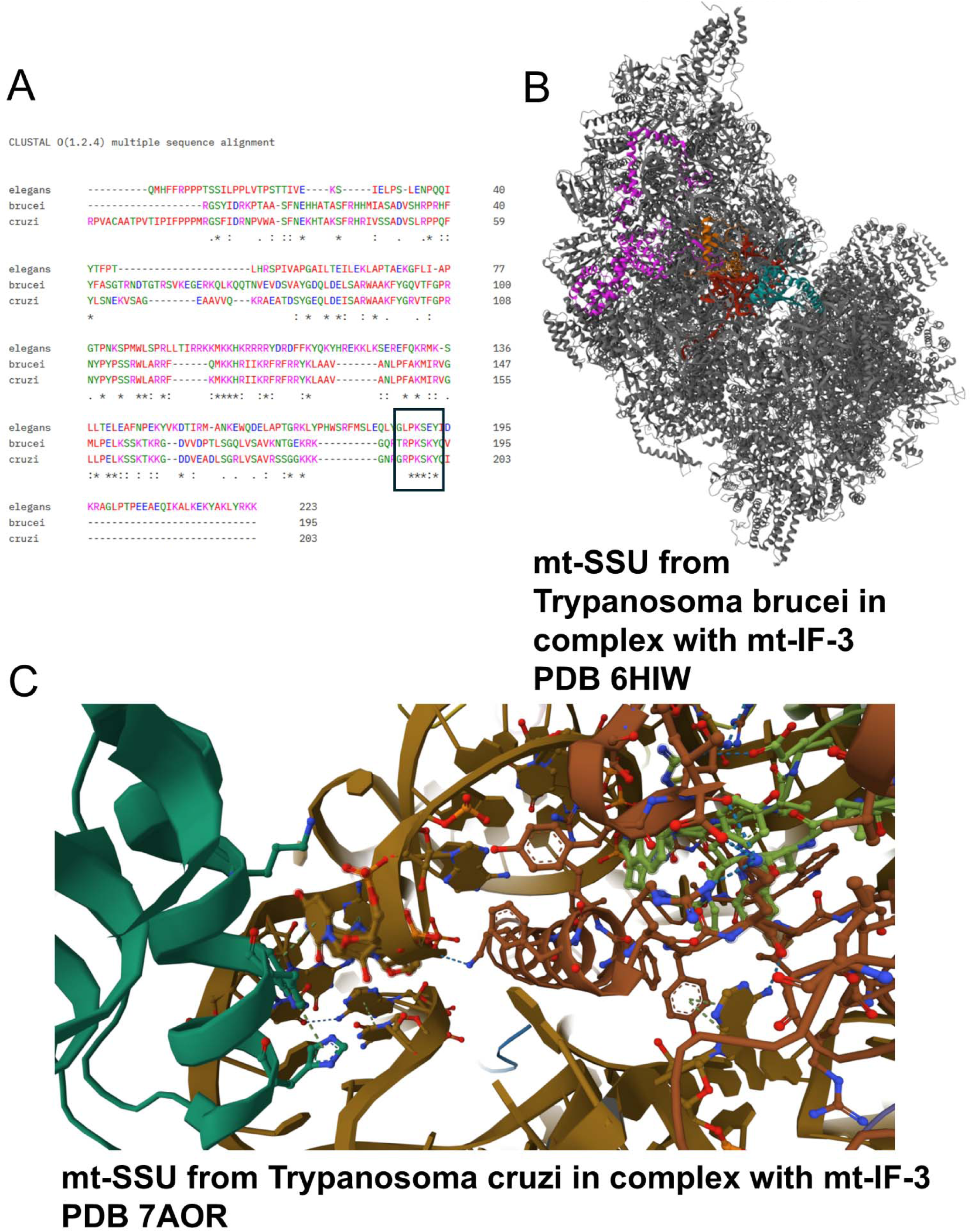
*mrps-38* homologs in *Trypanosma brucei* and *Trypanosoma cruzi* may regulate MTIF3. **(A)** ClustalO alignment of *C. elegans*, *T. brucei* and *T. cruzi* protein sequences, MRPS-38(P210L) region is outlined **(B)** Cryo-EM structure of Trypanosoma brucei small mitoribosomal subunit in complex with mt-IF-3 PDB: 6HIW^80^ MTIF3 is colored green, 9s rRNA is colored red, MRPS38 is colored orange and MRPS15 is colored purple. **(C)** Cryo-EM structure of Trypanosoma cruzi small mitoribosomal subunit in complex with mt-IF-3 PDB: 7AOR^82^ MTIF3 is colored dark green, 9s rRNA is colored gold, MRPS38 is colored brown and MRPS15 is colored light green.

**Figure S6.**
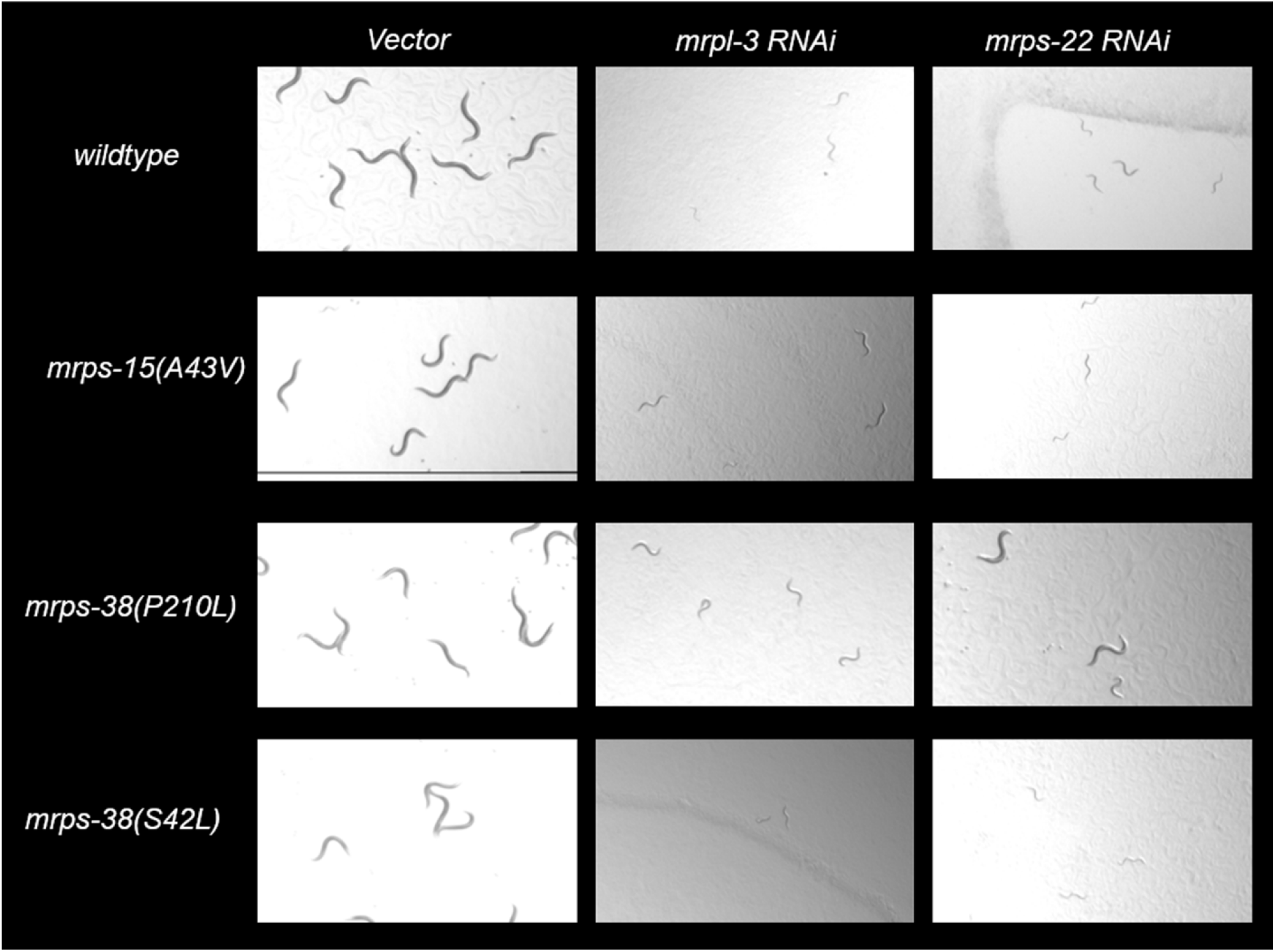
Knockdown of mrps-22 and mrpl-3 in mprs-15(A43V), mrps-38(P210L) and *mrps-38(S24L)* mutants. **(A)** Representative brightfield images of wildtype, *mrps-15(A43V), mrps-38(P210L) and mrps-38(S42L*) worms after 72 hours growth on *mrps-22* or *mrpl-3* RNAi at 11.3X magnification (scale bar, 1 mm). All depicted experiments had three biological replicates with similar results.

## Supplemental Tables

**Supplemental Table 1:** List of differentially expressed genes in *lonp-1(cmh24)* vs wildtype, *lonp-1(cmh24);mrps-38(P210L)* vs wildtype, *mrps-38(P210L)* vs wildtype and *lonp-1(cmh24)* vs *lonp-1(cmh24);mrps-38(P210L)* from RNA-seq.

**Supplemental Table 2:** Putative LONP-1 substrates from analysis of *C. elegans* mitochondrial proteomics^32–34^.

| Protein | *C. elegans* homolog | TMT Relative Abundance in wildtype | TMT Relative Abundance in *lonp-1(cmh24)* | TMT Relative Abundance in *lonp-1(cmh24); mrps-38(P210L)* | Putative substrate in *C. elegans*? | Reference |
| --- | --- | --- | --- | --- | --- | --- |
| ACO2 | ACO-2 | 6.28574 | 1.99762 | 10.62 | NO | 33 |
| MTERF D3 | ND | ND | ND | ND | ? | 33 |
| GAC | GLNA-1 | 4.22179 | 9.47785 | 7.08405 | YES | 33 |
| COX4 | COX-4 | 7.59094 | 2.82934 | 5.51736 | NO | 33 |
| PINK1 | PINK-1 | 5.00771 | 5.12891 | 5.10501 | NO | 33 |
| PARK7 | DJR-1.1 | 6.80904 | 8.10575 | 9.05951 | YES | 33 |
| ALAS-1 | ND | ND | ND | ND | ? | 33 |
| GLUT1 | ND | ND | ND | ND | ? | 33 |
| ERAL1 | ERAL-1 | 4.59307 | 9.39751 | 7.83498 | YES | 33 |
| SDHB | SDHB-1 | 8.20274 | 9.06651 | 7.44264 | NO | 33 |
| SDHAF2 | Y57A10A.29 | 2.49094 | 3.35606 | 6.21967 | YES | 33 |
| MRPL22 | T09F3.4 | 3.61988 | 5.80691 | 7.01845 | YES | 33 |
| CLPX | D2030.2 | 3.03255 | 5.40953 | 6.89577 | YES | 33 |
| STAR | STRL-1 | 4.7549 | 6.92371 | 5.65364 | YES | 33 |
| PMPCA | MPPA-1 | 3.93241 | 7.65569 | 7.72184 | YES | 33 |
| PMPCB | MPPB-1 | 1.30791 | 3.38213 | 8.83715 | YES | 33 |
| TFAM | HMG-5 | 4.0904 | 4.79578 | 6.54944 | YES | 33 |
| PDK4 | PDHK-2 | 3.77658 | 7.43631 | 9.46713 | YES | 33 |
| TWINKLE | TWNK-1 | ND | ND | ND | ? | 33 |
| RIP1 | ISP-1 | 6.96231 | 4.36467 | 5.70365 | NO | 32 |
| QCR2 | C05D11.1 | 8.15447 | 7.58781 | 8.29952 | NO | 32 |
| COX4 | COX5B | 5.57031 | 4.13309 | 8.56067 | NO | 32 |
| ATP1 | ATP-1 | 4.37464 | 7.66409 | 6.34474 | YES | 32 |
| ATP1 | T26E3.7 | 3.62926 | 1.94174 | 3.23488 | NO | 32 |
| ATP2 | ATP-2 | 7.51629 | 2.61273 | 6.46881 | NO | 32 |
| ATP7 | ATP-5 | 6.93031 | 3.2723 | 6.25565 | NO | 32 |
| DLD1 | F32D8.12 | 2.82731 | 2.77863 | 8.05301 | NO | 32 |
| Pdb1 | PDHB-1 | 5.34517 | 3.5696 | 6.79021 | NO | 32 |
| COQ5 | COQ-5 | 4.5356 | 5.23342 | 6.2352 | YES | 32 |
| ALD4 | ALH-3 | 3.08267 | 3.74907 | 3.86535 | YES | 32 |
| ALD4 | ALH-8 | 2.35777 | 1.93304 | 8.34757 | NO | 32 |
| ALD4 | ALH-9 | 9.33642 | 9.80145 | 10.9239 | YES | 32 |
| ALD4 | ALH-10 | 3.81179 | 4.13413 | 4.11773 | YES | 32 |
| ALD4 | ALH-11 | 5.46408 | 6.6679 | 7.75882 | YES | 32 |
| KGD2 | DLST-1 | 5.35695 | 2.65336 | 5.8281 | NO | 32 |
| MRP20 | MRPL-23 | 2.4993 | 2.90754 | 6.79694 | YES | 32 |
| PHB1 | PHB-1 | 5.88645 | 4.06312 | 6.14844 | NO | 32 |
| PHB2 | PHB-2 | 6.00642 | 6.52226 | 5.82346 | NO | 32 |
| AIM17 | GBH-1 | 4.00295 | 7.72243 | 5.2317 | YES | 32 |
| AIM17 | GBH-2 | 4.03749 | 6.58684 | 7.67244 | YES | 32 |
| LPD1 | DLD-1 | 5.76389 | 3.66767 | 6.51046 | NO | 32 |
| ILV5 | ND | ND | ND | ND | ? | 32 |
| HSP60 | HSP-60 | 5.4975 | 5.95936 | 5.30311 | NO | 32 |
| HSP78 | ND | ND | ND | ND | ? | 32 |
| SOD2 | SOD-2 | 6.30264 | 2.32978 | 10.9723 | NO | 32 |
| MRPL30 | MRPL-30 | 8.12557 | 8.00438 | 8.74231 | NO | 34 |
| MRPL1 | MRPL-1 | 6.21831 | 6.22342 | 6.05899 | NO | 34 |
| MRPL43 | MRPL-34 | 5.26835 | 3.61613 | 6.48537 | NO | 34 |
| MRPL11 | MRPL-11 | 6.3611 | 2.78624 | 6.4865 | NO | 34 |
| MRPL48 | MRPL-48 | 5.87432 | 3.48994 | 6.11144 | NO | 34 |
| MRPL21 | MRPL-21 | 8.20718 | 8.60715 | 8.2145 | NO | 34 |
| MRPS30 | MRPS-30 | 5.68393 | 8.04226 | 6.7466 | YES | 34 |
| MRPL44 | MRPL-44 | 5.43988 | 3.01711 | 6.02524 | NO | 34 |
| MRPL22 | MRPL-22 | 4.69119 | 9.04704 | 6.57631 | YES | 34 |
| FASTKD | FASK-1 | 4.31509 | 5.93022 | 6.69854 | YES | 34 |
| MRPS7 | MRPS-7 | 5.45415 | 4.82593 | 7.27032 | NO | 34 |
| MRPL55 | MRPL-55 | 2.2184 | 2.89136 | 6.30364 | NO | 34 |
| MRPS9 | MRPS-9 | 4.65927 | 4.03915 | 6.42493 | NO | 34 |
| MRPL3 | MRPS-18C | 3.04423 | 3.95044 | 7.04359 | YES | 34 |
| MRPL51 | MRPL-51 | 2.75459 | 3.82162 | 6.04768 | YES | 34 |
| POLRM<br>T | RPOM-1 | 7.03578 | 8.71289 | 8.36153 | YES | 34 |
| COA8 | Y39B6A.34 | 3.76496 | 4.53542 | 6.0778 | YES | 34 |
| MRPL46 | MRPL-46 | 1.6793 | 3.25298 | 7.81749 | YES | 34 |
| MRPS27 | MRPS-27 | 4.88454 | 6.78494 | 5.72555 | YES | 34 |
| DAP3 | DAP-3 | 2.63245 | 5.26486 | 7.6676 | YES | 34 |
| MRPL39 | MRPL-39 | 6.06618 | 6.85958 | 7.02082 | YES | 34 |
| MRPL49 | MRPL-49 | 4.82286 | 4.88631 | 6.83201 | YES | 34 |
| MRPL9 | MRPL-9 | 5.56704 | 7.31505 | 7.89861 | YES | 34 |
| PTCD3 | LET-630 | 3.56911 | 4.47856 | 7.39907 | YES | 34 |
| DDX28 | ND | ND | ND | ND | ? | 34 |
| MRPL37 | MRPL-37 | 2.5122 | 4.33701 | 6.86854 | YES | 34 |
| MRM1 | Y45F3A.9 | 2.99197 | 5.44679 | 6.6465 | YES | 34 |
| NUDT13 | NDX-9 | 3.56898 | 11.4966 | 10.4302 | YES | 34 |
| MRPS28 | MRPS-28 | 2.76746 | 4.06063 | 8.76371 | YES | 34 |
| MRPS14 | MRPS-14 | 2.52461 | 5.79465 | 8.02807 | YES | 34 |
| RBFA | C25G4.3 | 5.80875 | 9.70496 | 10.5466 | YES | 34 |
| LACTB2 | Y53F4B.39 | 6.88885 | 9.14456 | 6.81472 | NO | 34 |
| CLPX | K07A3.3 | ND | ND | ND | ? | 34 |
| CLPX | D2030.2 | 3.03255 | 5.40953 | 6.89577 | YES | 34 |
| GRSF1 | HRPF-2 | 4.54129 | 4.62731 | 5.18715 | YES | 34 |
| MRPL13 | Y53F4B.39 | 6.03003 | 8.02162 | 5.95886 | NO | 34 |
| TRUB2 | Y43B11AR.3 | 6.1681 | 9.44655 | 8.93233 | YES | 34 |
| MRPS6 | MRPS-6 | 2.79555 | 4.02665 | 6.80916 | YES | 34 |
| NDUFAF | NUAF-1 | 5.86196 | 9.04578 | 9.31974 | YES | 34 |
| CDK5R<br>A | RBM-12 | 4.04491 | 5.46256 | 8.54607 | YES | 34 |
| PDSS2 | COQ-1 | 3.14702 | 10.6859 | 5.48333 | YES | 34 |
| MTG1 | MTG-1 | 4.0863 | 9.16806 | 6.54898 | YES | 34 |
| METAP1 | MAP-1 | 4.96133 | 8.72978 | 7.70311 | YES | 34 |
| TRMT2B | TRM-2A | 5.73672 | 8.32942 | 5.07306 | NO | 34 |
| MRPS25 | MRPS-25 | 4.4191 | 4.25758 | 9.62217 | NO | 34 |
| MTRF1L | MTRF-1L | 5.78737 | 7.0066 | 8.76045 | YES | 34 |
| POLDIP | TAG-307 | 3.66904 | 10.8873 | 9.90782 | YES | 34 |
| MIPEP | MTG-1 | 4.17884 | 6.39455 | 8.76245 | YES | 34 |
| TFB2M | TFBM-1 | 7.42489 | 9.12467 | 8.06582 | YES | 34 |
| UQCC2 | Y48A5A.3 | 4.56767 | 4.8494 | 5.46114 | YES | 34 |
| PYCR2 | PYCR-1 | 5.03612 | 6.09159 | 4.93421 | NO | 34 |
| NSUN4 | NSUN-4 | 4.49344 | 5.65739 | 6.82479 | YES | 34 |
| MMADH<br>C | CELE_Y76A2<br>B.5 | 2.1415 | 10.6094 | 11.7018 | YES | 34 |
| MTPAP | C53A5.16 | 4.42205 | 6.28769 | 8.78208 | YES | This study |
| EARS2 | EARS-2 | 4.43746 | 9.89219 | 9.9764 | YES | This study |
| NARS2 | NARS-2 | 4.26453 | 7.87049 | 7.95794 | YES | This study |
| MTO1 | MTCU-2 | 3.02891 | 10.528 | 8.66918 | YES | This study |
| TCAIM | C35B8.3 | 3.63089 | 7.5044 | 7.40458 | YES | This study |
| MOCS1 | MOC-5 | 4.74118 | 9.76106 | 8.44878 | YES | This study |
| GTPBP3 | MTCU-1 | 3.82107 | 9.04768 | 8.48203 | YES | This study |
| SLIRP | SLRP-1 | 1.67713 | 10.5175 | 11.1674 | YES | This study |
| EARS2 | EARS-2 | 4.43746 | 9.89219 | 9.9764 | YES | This study |
| QRSL1 | Y41D4A.6 | 5.29036 | 10.5415 | 11.3769 | YES | This study |
| POLG | POLG-1 | 4.15084 | 11.2368 | 11.4913 | YES | This study |
| ATF5 | ATFS-1 | 1.92776 | 14.6991 | 14.0918 | YES | This study |
| RBFA | C25G4.3 | 5.80875 | 9.70496 | 10.5466 | YES | This study |
| DARS2 | DARS-2 | 5.95267 | 11.214 | 9.27879 | YES | This study |
| TRMU | MTTU-1 | 2.94315 | 14.018 | 10.2419 | YES | This study |
| C1ORF5<br>3 | MRPL-14 | 4.09998 | 10.2499 | 10.1241 | YES | This study |
| NDUFAF<br>1 | NUAF-1 | 5.86196 | 9.04578 | 9.31974 | YES | This study |
| NOA1 | NOA-1 | 6.72617 | 9.12998 | 8.85212 | YES | This study |
| GFM2 | Y119D3B.14 | 3.81019 | 6.51167 | 7.47824 | YES | This study |
| MRPS11 | MRPS-11 | 2.60413 | 4.88019 | 8.20778 | YES | This study |
| CLPP | CLPP-1 | 8.1006 | 11.1718 | 9.70155 | YES | This study |

**Supplemental Table 3:** Table of putative *C. elegans* mitochondrial proteins from Li et al.^27^, Lesnik et al.^36^ and Kenny-Ganzert et al^37^.

**Supplemental Table 4:** List of differentially expressed proteins in *lonp-1(cmh24)* vs wildtype, *lonp-1(cmh24);mrps-38(P210L)* vs wildtype, *mrps-38(P210L)* vs wildtype and *lonp-1(cmh24)* vs *lonp-1(cmh24);mrps-38(P210L)* from mass spectrometry.

**Supplemental Table 5:** *C. elegans* homologs of human OXPHOS proteins.

| Protein | *C. elegans* homolog | Complex |
| --- | --- | --- |
| ND1 | NDUO-1 | I |
| ND 2 | NDUO-2 | I |
| ND 4 | NDUO-4 | I |
| ND 5 | NDUO-5 | I |
| NDUFA1 | F31D4.9 | I |
| NDUFA2 | Y63D3A.7 | I |
| NDUFA5 | C33A12.1 | I |
| NDUFA6 | NUO-3 | I |
| NDUFA7 | F45H10.3 | I |
| NDUFA8 | Y54F10AM.5 | I |
| NDUFA9 | NDUF-9 | I |
| NDUFA10 | NUO-4 | I |
| NDUFA11 | NDUF-11 | I |
| NDUFA12 | Y94H6A.8 | I |
| NDUFAB1-2 | Y56A3A.19 | I |
| NDUFA13 | C34B2.8 | I |
| NDUFB2 | F44G4.2 | I |
| NDUFB3 | C18E9.4 | I |
| NDUFB4 | NUO-6 | I |
| NDUFB5 | C25H3.9 | I |
| NDUFB6 | ZK809.3 | I |
| NDUFB7 | D2030.4 | I |
| NDUFB8 | Y51H1A.3 | I |
| NDUFB9 | C16A3.5 | I |
| NDUFB10 | F59C6.5 | I |
| NDUFB11 | F42G8.10 | I |
| NDUFC2 | Y71H2AM.4 | I |
| NDUFS1 | NUO-5 | I |
| NDUFS2 | GAS-1 | I |
| NDUFS3 | NUO-2 | I |
| NDUFS4 | LPD-5 | I |
| NDUFS5 | NDUF-5 | I |
| NDUFS6 | NDUF-6 | I |
| NDUFS7 | NDUF-7 | I |
| NDUFS8 | T20H4.5 | I |
| NDUFV1 | NUO-1 | I |
| NDUFV2 | F53F4.10 | I |
| NDUFV3 | Y69A2AR.3 | I |
| SDHA | SDHA-1 | II |
| SDHA | SDHA-2 | II |
| SDHB | SDHB-1 | II |
| SHDC | MEV-1 | II |
| SDHD | SDHD-1 | II |
| MT CTB | CTB-1 | III |
| UQCRFS1 | ISP-1 | III |
| UQCRB | T02H6.11 | III |
| UQCRC1 | UCR-1 | III |
| UQCRC2 | UCR-2.1 | III |
| UQCRC2 | UCR-2.2 | III |
| UQCRC2 | UCR-2.3 | III |
| UQCRH | T27E9.2 | III |
| UQCRQ | R07E4.3 | III |
| UQCRQ | F45H10.2 | III |
| UQCR10 | C14B9.10 | III |
| UQCR11 | UCR-11 | III |
| CYC1 | CYC-1 | III |
| UQCRHL | T27E9.2 | III |
| MT-COI | CTC-1 | IV |
| MT-COII | CTC-2 | IV |
| COX4I1 | COX-4 | IV |
| COX5A | COX-5A | IV |
| COX5B | COX-5B | IV |
| COX6A1 | COX-6A | IV |
| COX6B1 | COX-6B | IV |
| COX6C | COX-6C | IV |
| COX7C | COX-7C | IV |
| MT-ATP6 | ATP-6 | V |
| ATP5F1A | ATP-1 | V |
| ATP5F1B | ATP-2 | V |
| ATP5F1D | F58F12.1 | V |
| ATP5F1E | HPO-18 | V |
| ATP5F1E | ZC262.5 | V |
| ATP5F1G | Y69A2AR.18 | V |
| ATP5J2 | R53.4 | V |
| ATP5MC2 | Y82E9BR.3 | V |
| ATP5ME | R04F11.2 | V |
| ATP5MG | ASG-1 | V |
| ATP5MG | ASG-2 | V |
| ATP5PB | ASB-1 | V |
| ATP5PB | ASB-2 | V |
| ATP5PD | ATP-5 | V |
| ATP5PF | ATP-4 | V |
| ATP5PO | ATP-3 | V |
| ATP5SL | Y53G8AR.8 | V |
| ATP5MF | R53.4 | V |

**Supplemental Table 6:** Gene Ontology results from STRING analysis of the Top 100 CRISPR Co-dependencies of human LONP1 from DepMap^56,74^.

